# Event structure sculpts neural population dynamics in the lateral entorhinal cortex

**DOI:** 10.1101/2024.06.17.599402

**Authors:** Benjamin R. Kanter, Christine M. Lykken, May-Britt Moser, Edvard I. Moser

## Abstract

Our experience of the world is a continuous stream of events which must be segmented and organized simultaneously at multiple timescales. The neural mechanisms underlying this process remain unknown. Here, we simultaneously recorded many hundreds of neurons in the lateral entorhinal cortex (LEC) of freely behaving rats as we manipulated event structure at multiple timescales. During foraging as well as during sleep, population activity drifted continuously and unidirectionally along a one-dimensional manifold. Boundaries between events were associated with discrete shifts in state space, suggesting that LEC dynamics directly reflect event segmentation. During tasks with a recurring temporal structure, activity traveled additionally in directions orthogonal to the flow of drift, enabling the LEC population to multiplex event information across different timescales. Taken together, these results identify a hierarchically organized neural coding scheme for segmenting and organizing events in time.

---

We experience the world as a continuous stream of events (*1*), consisting of things occurring in a particular order at particular places and times. Episodic memory allows us to mentally revisit those experiences by recalling events in sequence (*2*). While the hippocampus is critical for episodic memory (*3, 4*), and the spatial correlates of such memories have been well described (*5, 6*), much less is known about the neural mechanisms underlying the temporal organization of episodic memories (*7*). The passage of time is mirrored by slow drift in neural activity in the hippocampus (*8–12*) and one of its major cortical inputs, the lateral entorhinal cortex (LEC) (*13, 14*), but the contribution of this drift to the temporal organization of episodic memories has not been determined. In particular, it remains unknown whether the drift of neural population activity is steady and continuous in time, whether it reflects the structure of experience, and whether and how experience is encoded simultaneously at multiple timescales (*15, 16*).

One important clue is that experience is hierarchically segmented into discrete events across a range of timescales from seconds to minutes or more (*17*). Event boundaries (i.e., times of transition between events) are associated with abrupt changes in the external environment or in physical location. Such boundaries have profound effects on memory for the duration and order of events (*9, 17–21*) and are accompanied by transient changes in hippocampal activity (*16, 21–24*). To search for the neural mechanisms that determine how events are segmented and organized in time, we monitored the activity of large populations of individually separable neurons during single episodes of experience in brain areas where neural activity is correlated with the passage of time. We took advantage of newly developed high-density Neuropixels 2.0 silicon probes (*25*) to perform simultaneous extracellular recordings of more than one thousand neurons in LEC, medial entorhinal cortex (MEC), and hippocampal area CA1 (CA1) of freely behaving rats as we manipulated event structure at multiple timescales.

## Continuous one-dimensional drift of LEC population activity

We first sought to characterize LEC dynamics during unconstrained behavior in the absence of scheduled events using a free foraging task where rats explored an open field arena and chased food crumbs thrown in random locations at variable intervals by the experimenter. We used a similar paradigm previously to observe that LEC population activity drifts over time (*13*), but by increasing cell yield by an order of magnitude we were now able to avoid pooling neurons across experiments and could instead directly quantify neural dynamics during individual events, a necessary condition for relating activity to the structure of experience (Fig. 1A; mean = 494 LEC neurons per session, 26 sessions, 7 rats; table S1, fig. S1-2). To visualize LEC dynamics over the course of the session, we first temporally binned the spiking activity of each LEC neuron in 10- sec bins. Next, we ran principal component analysis (PCA) on this unsmoothed spike count matrix (time × neurons) and kept the components that explained 50% of the variance to moderately denoise the activity. Lastly, we ran linear discriminant analysis using clock time as class labels (1-min epochs) to find dimensions along which the population activity varied most over time.

**Fig. 1.**
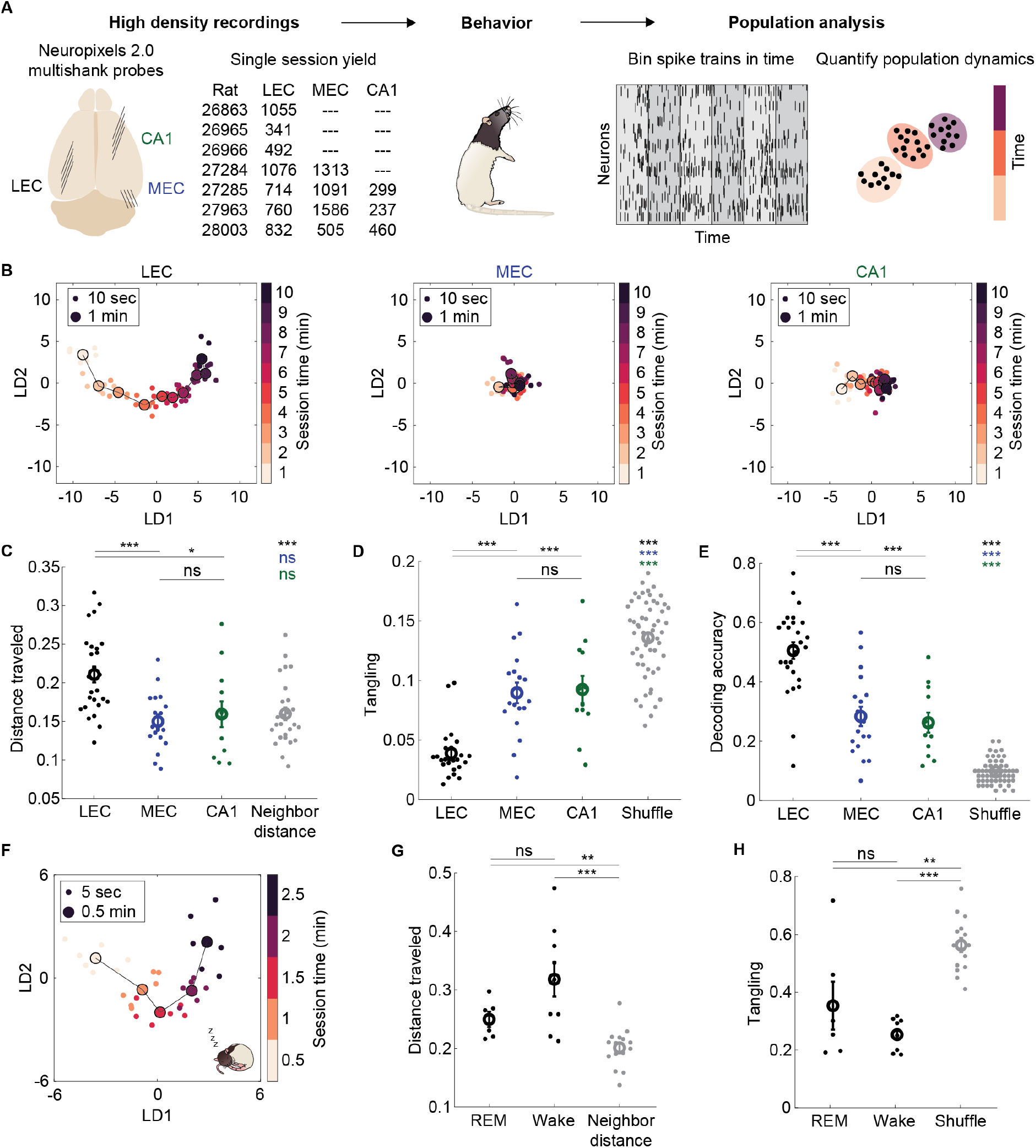
LEC population activity continuously drifts along a one-dimensional manifold without reversing direction. (**A**) Summary of experimental approach. Neuropixels 2.0 silicon probes were used to record large populations of neurons simultaneously in one to three brain areas. Rats performed behavioral tasks with different degrees of event structure. Population dynamics were visualized and quantified during individual experiences. (**B**) Example state space trajectories (from rat 27285) during 10 min of free foraging show population drift in LEC (left) that is largely absent in simultaneously recorded MEC (middle) and CA1 (right) populations. State space is defined by top two linear discriminants LD1 and LD2 using 1-min epochs as class labels (see Methods). Small dots represent 10-sec time bins, large dots represent average activity during 1-min epochs. Points are colored from light to dark to show time within the session. (**C**) Distance traveled between first and last minutes for all sessions and all areas, calculated as cosine distance between population vectors in the full n-dimensional state space. Neighbor distances compare adjacent times in LEC as a lower bound of distance traveled. LEC vs MEC: t(42) = 4.24, p = 1.19e^-4^, LEC vs CA1: t(36) = 2.72, p = 0.01, MEC vs CA1: t(28) = -0.55, p = 0.59, LEC vs neighbor: t(50) = 3.86, p = 3.25e^-4^, MEC vs neighbor: t(42) = -0.81, p = 0.43, CA1 vs neighbor: t(36) = -0.03, p = 0.98, two-sample t-test; n = 26, 18, and 12 sessions for LEC, MEC, and CA1, respectively. (**D**) Tangling of neural trajectories quantifies the extent to which nearby state space locations have different movement directions. Tangling was calculated in the 2D subspace (LD1 vs LD2). Shuffle obtained by shuffling epoch labels. LEC vs MEC: t(42) = -5.95, p = 4.66e^-7^, LEC vs CA1: t(36) = -5.63, p = 2.16e^-6^, MEC vs CA1: t(28) = -0.21, p = 0.84, LEC vs shuffle: t(80) = -14.14, p = 1.84e^-23^, MEC vs shuffle : t(72) = -5.13, p = 2.32e^-6^, CA1 vs shuffle : t(66) = - 4.07, p = 1.27e^-4^, two-sample t-test; n = 26, 18, and 12 sessions for LEC, MEC, and CA1, respectively. (**E**) Decoding accuracy for 1-min epochs within the session using a linear classifier. Decoding performed on principal components explaining 50% of variance. LEC vs MEC: t(42) = 5.13, p = 6.92e^-6^, LEC vs CA1: t(36) = 5.14, p = 9.83e^-6^, MEC vs CA1: t(28) = 0.43, p = 0.67, LEC vs shuffle: t(80) = 19.90, p = 1.03e^-32^, MEC vs shuffle: t(72) = 9.05, p = 1.71e^-13^, CA1 vs shuffle: t(66) = 8.64, p = 1.94e^-12^, two-sample t-test; n = 26, 18, and 12 sessions for LEC, MEC, and CA1, respectively. (**F**) Trajectory in LEC during an example 2.5-min bout of REM sleep. (**G**) Distance traveled between first and last time bins in REM sleep compared to duration-matched data during wake in LEC. REM vs wake: t(13) = -1.80, p =0.10, REM vs neighbor: t(19) = 3.04, p = 6.70e^-3^, wake vs neighbor: t(22) = 4.65, p = 1.23e^-4^, two-sample t-test; n = 6 and 9 bouts for REM and wake, respectively. (**H**) Tangling in REM sleep compared to duration-matched data during wake. REM vs wake: t(13) = 1.43, p =0.18, REM vs neighbor: t(19) = -3.28, p = 3.90e^-3^, wake vs neighbor: t(22) = -8.97, p = 8.32e^-9^, two-sample t-test; n = 6 and 9 bouts for REM and wake, respectively. **(B-H**) Data represented as individual foraging sessions (or REM bouts) and mean +/- SEM. ***p < 0.001, **p < 0.01, *p < 0.05, ns = not significant.

We first replicated our previous work (*13*) showing that LEC population activity drifted over the course of minutes (Fig. 1B, left; fig. S3-4). This drift creates trajectories in state space that are ordered in time such that the population activity at any given time is most similar to the neighboring timepoints and becomes more dissimilar as time evolves (quantified below). There was little or no drift in neighboring MEC or CA1 during the exact same experience (Fig. 1B, middle and right; fig. S3-4). To quantify the amount of drift in LEC and to directly compare it to MEC and CA1, we calculated the distance traveled in state space during the 10-min foraging session. Distance traveled from the first to last minute of the 10-min session was defined as the cosine distance between population vectors, calculated in the full n-dimensional space (n = neurons) to avoid potential distortions introduced by dimensionality reduction. LEC activity drifted significantly farther than MEC and CA1 during the 10 min of foraging (Fig. 1C; fig. S4C). Distance travelled from the first minute increased progressively during the session (Fig. 1B and 1C; fig. S3).

The large population recordings achieved with Neuropixels probes enabled us to determine the dynamics of neural trajectories during unique experiences. By simultaneously recording trajectories in LEC, MEC, and CA1, we were able to show that LEC drifts more than the other regions during the exact same experience (fig. S4C). We then quantified whether these trajectories continuously visited new positions in state space over time, or whether they might loop back to the same positions again. Such tangled trajectories would not provide an accurate measure of time within an episode because the same neural activity would be associated with multiple timepoints. We calculated the degree of trajectory tangling in the state space defined by the top two linear discriminants in the following way: for any pair of points along the trajectory, we compared the difference in their derivatives to their distance in state space. Intuitively, if points close together have different derivatives there is high tangling, whereas if they have the same derivative tangling approaches zero. LEC trajectories had minimal tangling compared to MEC and CA1 (Fig. 1D), consistent with LEC activity evolving along a one-dimensional manifold. To rule out that trajectories followed the same path multiple times (e.g., traveling around a ring-shaped manifold would yield minimal tangling), we decoded time within the foraging session based on activity from either LEC, MEC, or CA1. We found that a linear decoder was sufficient to decode time from LEC activity with an accuracy twice as high as MEC and CA1, and over five times higher than chance (Fig. 1E; fig. S4B). This result confirms that LEC activity evolves unidirectionally such that each timepoint generates a unique population state. Downstream brain areas can therefore linearly read out temporal information from LEC population vectors.

After demonstrating that LEC population activity drifted in the absence of scheduled events, we next asked whether it also drifted in the absence of sensory inputs that normally guide behavior. We recorded LEC activity as rats voluntarily slept in a small box and focused on epochs of rapid eye movement (REM) sleep, when neural dynamics resemble wakefulness. Neural trajectories were comparable to those found during foraging (Fig. 1F). To quantify the drift and make a fair comparison between REM and wake, we selected REM epochs lasting 2.5 min and compared them to 2.5-min bouts of foraging data from the same rats. The amount of drift during REM was not different from wake (Fig. 1G), and there was no difference in the amount of tangling (Fig. 1H). These results imply that continuous drift is an inherent property of LEC population activity rather than one that requires ongoing experience. This is consistent with observations of drift already during the animals’ first experience in the foraging task (Fig. S4D).

## Shifts in state space segment events

After showing that LEC dynamics evolve continuously in time without relying on transitions between events to propel the drift, we next asked whether and how event boundaries may modulate the inherently generated drift. One clue is the fact that LEC neurons encode stimuli such as odors and objects via time-locked changes in firing rate (*26–30*). At the population level, such abrupt changes in firing rates could evoke abrupt shifts in state space, which would be detectable as brief moments of acceleration then deceleration of the neural trajectory (Fig. 2A). This would provide a simple mechanism for ensembles of co-active neurons to timestamp event boundaries.

**Fig. 2.**
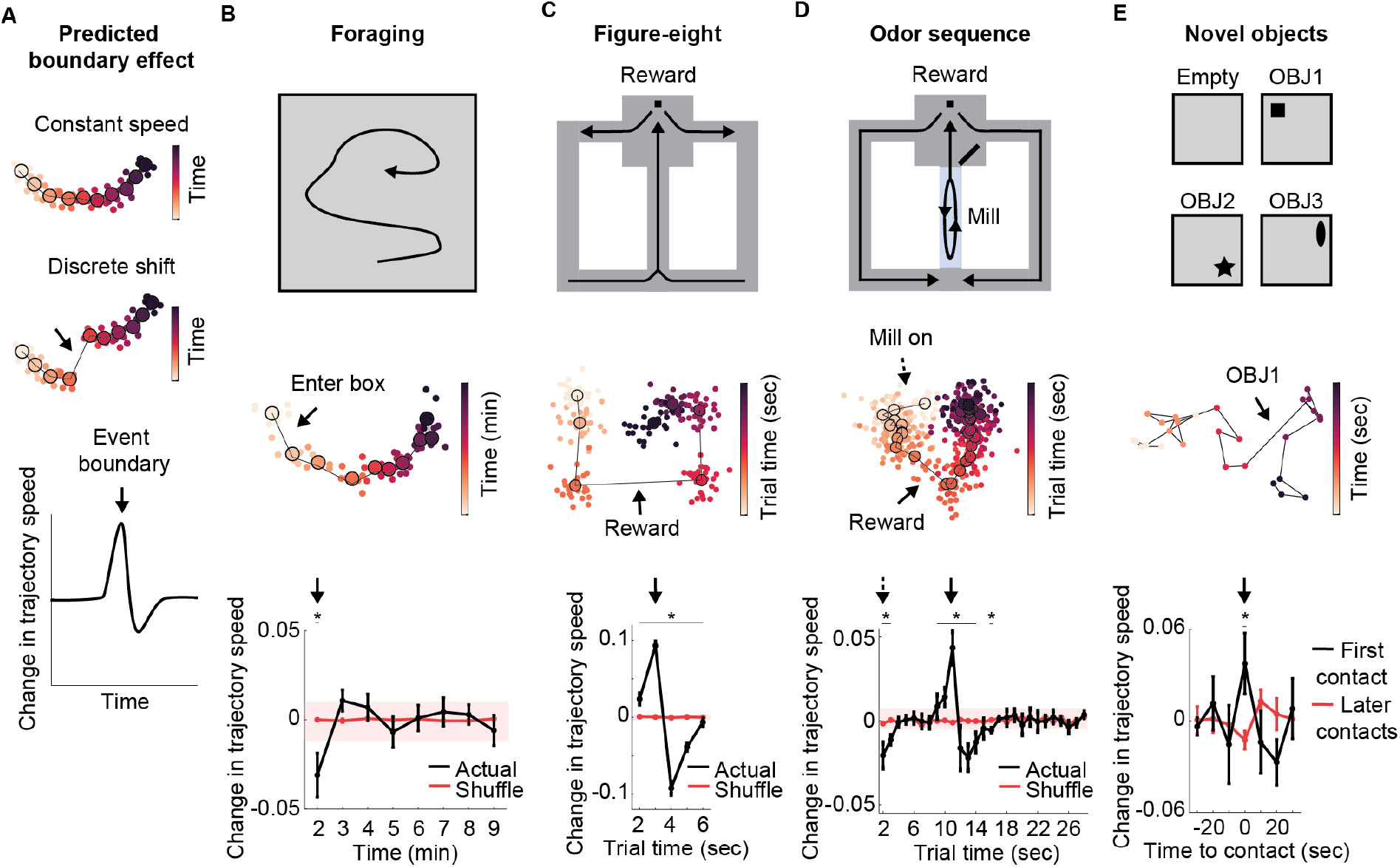
Shifts in state space at event boundaries discretize experience. (**A**) Schematic illustrating the hypothesis that trajectories evolve at a constant speed within an event (top) and undergo discrete shifts at event boundaries (middle). Plotting the change in trajectory in speed over time reveals a rapid acceleration then deceleration at the event boundary (bottom). (**B**) In the foraging task, entering the box was the only scheduled event boundary (top). Neural trajectories decelerated after entering the box and then returned to a constant speed. Example trajectory (middle) with arrow indicating change in speed. Mean change in trajectory speed (bottom) at each timepoint for all sessions compared to control obtained by shuffling timepoints. Stars mark timepoints where mean is below/above the 1^st^/99^th^ percentile of the shuffled distribution (shaded region). Interaction effect between group and time: F(7, 336) = 2.3, p = 0.03, repeated measures ANOVA, n = 26 sessions. (**C**) In the figure-eight task, the reward on each trial (lap) was the only scheduled event boundary. Neural trajectories accelerated during reward approach and then decelerated again on each trial. Data displayed as in B. Interaction effect between group and time: F(4, 32) = 142.5, p << 0.001, repeated measures ANOVA, n = 5 sessions. (**D**) In the odor sequence task, there were two scheduled event boundaries per trial. Neural trajectories decelerated after the treadmill turned on to start the trial. Neural trajectories accelerated during reward approach and then decelerated again. Data displayed as in B. Interaction effect between group and time: F(26, 156) = 5.0, p = 6.75e^-11^, repeated measures ANOVA, n = 4 sessions. (**E**) In the novel objects task, the objects defined the event boundaries. Neural trajectories accelerated at the first contact with each object, but not for subsequent contacts with the same object. Example trajectory (middle) during contact with OBJ1. Dots represent 10-sec time bins, colors represent 1-min epochs. Mean change in trajectory speed (bottom) for first contacts (black) versus later contacts (red). t(33) = 3.16, p = 3.40e^-3^, two-sample t-test, n = 35 timepoints. t-test instead of ANOVA as in other panels due to the limited number of first contacts. Star here marks significance after Bonferroni correction for multiple comparisons at α=0.05. **(B-D)** Shuffle obtained by shuffling timepoints. (**B-E**) Data represented as mean +/- SEM. Change in trajectory speed calculated as the difference between cosine distance in the full n-dimensional space (i.e., population vectors) for neighboring timepoints and the previous two neighboring timepoints.

We searched for such signatures of event segmentation in LEC by measuring the acceleration profiles of neural trajectories in the full n-dimensional state space in tasks with different types of events over multiple timescales. In the foraging task described above, there were no scheduled event boundaries aside from the introduction of the animal to the arena (animals could not predict when the session would end because sessions were of variable duration and truncated post-hoc to 10 min for analysis) (Fig. 2B). We found a deceleration of trajectory speed during the first minutes of the session, consistent with a post-boundary deceleration to baseline speed. Next, rats were trained to run self-paced laps around a figure-eight maze, motivated by a single reward of sweetened chocolate milk at a constant location on each lap (Fig. 2C). Trajectories accelerated on each lap immediately before the animal reached the reward, which was the one scheduled event boundary in the task, and then decelerated again. Lastly, we used an odor sequence task that was similar in many ways to the figure-eight task. The only relevant difference here is that the central stem of the maze was a treadmill where the rat ran in place for 10 sec, creating an additional event within each trial (other task features described below). We observed multiple changes in trajectory acceleration aligned to the multiple event boundaries within each trial of this task (Fig. 2D): a deceleration after the treadmill turned on, an acceleration during reward approach, and a deceleration upon reaching the reward. Note how stable the trajectory speeds were within each event compared to the boundary-induced shifts. Together, these results indicate that shifts in LEC activity are consistently time-locked to event boundaries across a variety of behaviors.

To test whether LEC is also sensitive to novel event boundaries, prior to learning, we next scheduled event boundaries at times when neural trajectory speeds were known to be stable. Animals started by randomly foraging in an empty arena, as above, but every 7.5 min a novel object was inserted at a pseudorandom location (Fig. 2E, top). We predicted that object exploration would elicit shifts in LEC activity due to the coincident activation of object- responsive neurons (*26, 27*). Indeed, we found that times of object exploration caused higher firing rates in LEC neurons compared to all other time points (fig. S5A). Importantly, the first exploration of each object was associated with acceleration of the neural trajectory at the time of contact with the object (Fig. 2E), similar to the familiar event boundaries in the other tasks. Subsequent exploration of the same objects did not cause shifts (Fig. 2E). To avoid biasing our search for shifts to periods of object exploration, we also performed an agnostic search of the whole experiment for any times where the network became suddenly active as putative times for shifts. We identified such times as moments with the population rate exceeded its 90^th^ percentile and the increase in population rate from the previous time bin also exceeded its 90^th^ percentile. These onsets of synchronous activity were consistently associated with trajectory acceleration (fig. S5B). The majority of these synchronous times corresponded to moments of object exploration (9/15 or 60% within 10 cm radius of object). Similar results were obtained for agnostic searches across the trial-based tasks (figure-eight task: 62% +-/ 8% near reward location, n = 5 sessions; odor sequence task: 92% +/- 2% near reward locations or treadmill start, n = 4 sessions). We thus conclude that event boundaries defined externally as salient changes in the world or internally as shifts in neural state space are two sides of the same coin (*23*).

## Multiplexing timescales via orthogonal coding dimensions

Event memories are structured at multiple timescales, from seconds to minutes or hours. We asked whether different timescales could be encoded simultaneously in LEC. The trial-based tasks described above contain a hierarchy of timescales with individual laps occurring over seconds and the behavioral session occurring over minutes. To determine if LEC contains representations matching each of these timescales during ongoing behavior, we trained rats in a repetitive lap running task that caused repetitive neural trajectories in LEC (*13*) (Fig. 3A; fig. S6A-B). From the continuous behavior, we extracted 6-sec trials leading up to the reward. As expected from our previous work (*13*), LEC activity followed similar trajectories during these 6 sec on each lap (trial time; Fig. 3B,E; fig. S6C-E). The large population recordings of the present study enabled us to analyze individual trials and compare the trajectories lap-by-lap. We directly quantified the alignment of the trajectories by calculating the distances between matched trial times across trials and comparing them to the distances between mismatched trial times (Fig. 3C; cosine distances in full state space). Subtracting matched distances from mismatched distances gives an alignment score where positive values indicate that matched times are closer together in state space (i.e., the trial trajectories are in register). Trajectories were significantly better aligned across trials compared to data where trial times were shuffled (Fig. 3C), and alignments were maintained between trajectories for different trial types (left versus right turn trials) (fig. S6E). We further confirmed that activity repeated across trials by accurately decoding trial time from held out trials using a linear decoder (Fig. 3D). Altogether, these observations suggest that LEC activity followed stereotypic trajectories over the course of each trial, during the seconds leading up to reward.

**Fig. 3.**
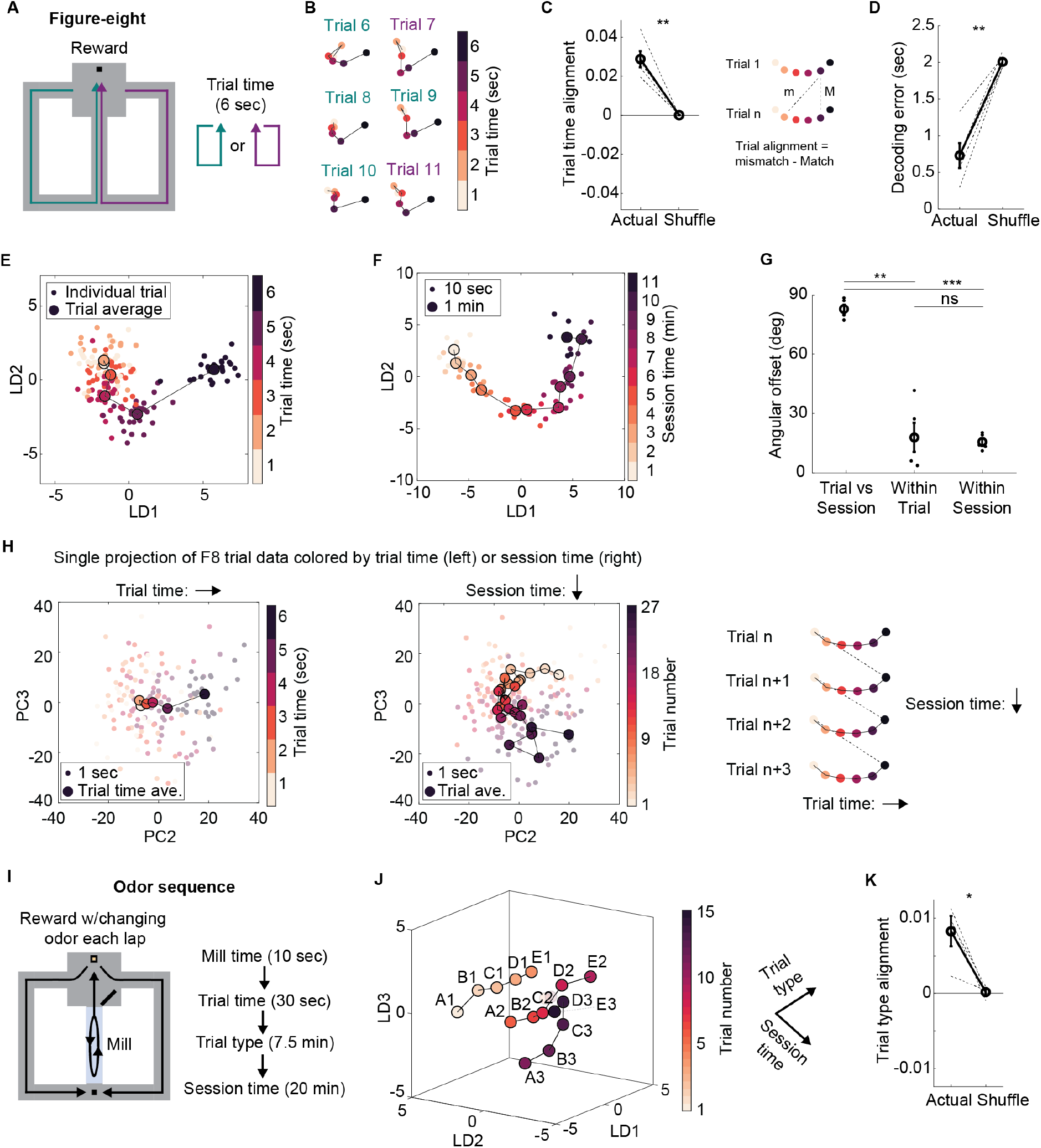
Diverse timescales are multiplexed in LEC activity using orthogonal coding dimensions. (**A**) In the figure-eight task, rats were trained to run self-paced laps to receive milk rewards at the top of the maze, returning down the arm of their choice. Trial times were defined post-hoc as 6 sec leading up to but excluding the reward. (**B**) Example trajectories from light to dark points during individual trials show that trajectories were aligned across trials. (**C**) At each trial time, activity was closer to matched trial times from other trials compared to mismatched times from other trials (left). Actual vs shuffle: t(4) = 6.85, p = 2.40e-3, paired t-test, n = 5 sessions. Schematic (right) shows how alignment was calculated as cosine distance in full dimensional space between mismatched trial times minus matched trial times across all trial pairs. (**D**) Decoding accuracy for 1-sec trial times using a linear classifier trained on held out trials. Actual vs shuffle: t(4) = -8.41, p = 1.11e^-3^, paired t-test; n = 5 sessions. (**E**) Trial-averaged trajectory for example session where small dots represent 1-sec time bins on each trial, large dots represent average activity for that time bin over all trials. Points are colored from light to dark to show time within the trial. (**F**) Ignoring trial structure and looking for drift as in the foraging task shows that repeating trajectories across trials did not eliminate drift over the course of the session. Small dots represent 10-sec time bins, large dots represent average activity for 1-min epochs. Points are colored from light to dark to show time within the session. (**G**) The axes of travel during each trial vs during the session were orthogonal. Trial-Session vs Within trial: t(4) = 8.22, p = 1.20e-3, Trial-Session vs Within session: t(4) = 23.68, p = 1.89e-5, Within trial vs Within session: t(4) = 0.28, p = 0.80; paired t-test, n = 5 sessions. (**H**) Same example session from (E-F) showing a single projection of trial data in the subspace defined by PC2 and PC3. The data is colored by either trial time (left) or session time (middle) to further validate that these different timescales are represented by orthogonal axes. Schematic (right) shows how activity can repeat across trials while simultaneously drifting in an orthogonal direction. (**I**) In the odor sequence task, rats were trained to run self-paced laps starting with a 10-sec treadmill period, then retrieving a buried chocolate reward in an odorized cup of sand, and finally returning to the base of the maze for a small milk reward. Each lap constituted one trial. Five odors were presented on consecutive laps forming a sequence that was followed by a 5-min rest off of the maze. Each session contained three sequence runs, yielding a total of fifteen laps. Schematic (right) shows hierarchy of timescales from a few seconds to many minutes. (**J**) Example session showing sequence runs traced parallel trajectories through state space, which could serve to link temporal contexts. Dots represent average activity for each trial. Points are colored from light to dark to show time within the session. Trial type and session time were approximately orthogonal. (**K**) Activity was more similar for matched trial types compared to mismatched trial types across sequence runs. Actual vs shuffle: t(3) = 3.79, p = 0.03, paired t-test, n = 4 sessions. (**C, D, G, K**) Data represented as mean +/- SEM. ***p < 0.001, **p < 0.01, *p < 0.05, ns = not significant.

This finding of repeating trajectories during repetitive behavior is quite the opposite of continuous drift. To determine if the repetitive nature of the task impacted dynamics also at slower timescales, we next zoomed out to the timescale of the behavioral session and asked whether drift over minutes was preserved. We performed the exact same analysis as in the foraging task to capture change over minutes while ignoring the lap running behavior. Continuous drift was still observed (Fig. 3F, fig. S6F-G) during the exact same behavioral session while repeating trajectories occurred at a faster timescale (Fig. 3E). This suggests that drift and repeating trajectories toward learned event boundaries may evolve along independent dimensions in state space and that information about these two timescales are multiplexed (fig. S6H-I) within the same neural population.

To quantify whether these two coding dimensions – session time and trial time – were orthogonal, we calculated the angle between them in the full state space. We found the angle between coding dimensions to be approximately 90 degrees, which was significantly larger than the variability within each coding dimension (ca. 15 degrees) calculated using a resampling procedure (Fig. 3G; see Methods). However, while this approach was well-suited to identifying the best angle for each timescale, it required two separate sources of input data (trial-based data with 1-sec bins vs full session data with 10-sec bins), which prevented us from observing the orthogonality in a single subspace. We therefore used a complementary approach, running PCA on the trial-based data with 1-sec bins and asking whether any of the top principal components were well correlated to either trial time or session time. In some cases, there were strong correlations to both trial and session time such that we could visualize both coding dimensions in a single 2D subspace defined by those principal components (Fig. 3H, left, middle), which are orthogonal by definition. Combining these two orthogonal coding dimensions yields a helical trajectory where each coil of the helix represents the recurring activity for each trial and the long axis of the helix represents continuous drift throughout the session (Fig. 3H, right).

Real world experiences, however, do not contain a single recurring event (e.g., one reward per trial), but rather consist of many different events across diverse timescales. To test whether LEC activity could evolve simultaneously along a larger number of trajectories, we used an odor sequence task (briefly mentioned above) with recurring, hierarchically organized events spanning timescales of seconds to many minutes (Fig. 3I). The rat first ran in place on a treadmill for 10 sec. Next, they ran one lap around the figure-eight maze stopping to sample an odorized cup of sand and dig for a buried chocolate reward. The odor changed on each lap such that the odors formed a sequence over five laps from odor A to odor E. Lastly, they performed three sequence runs (i.e., 15 total trials) with a 5-min rest between runs.

At the short timescale of seconds, LEC activity exhibited repeating trajectories during each trial. LEC activity was more similar between trials for matched trial times compared to mismatched trial times relative to shuffled controls (fig. S7A). These trial-based trajectories were similar to those in the figure-eight task above, but extended for approximately 30 sec, demonstrating that LEC activity can capture the fine temporal details of extended experience. Moreover, trajectories were also aligned across repeated periods on the treadmill (fig. S7B) in the absence of overt changes in the external environment. Time on the mill could be accurately decoded from held out trials using a linear decoder (fig. S7C).

At the long timescale of minutes, LEC activity drifted as in all tasks described above (fig. S7D). The activity evolved smoothly from one trial to the next along a linear trajectory during the first sequence run. After a 5-min rest, the activity did not continue where it left off. It also did not retrace the same trajectory or reset and evolve in an arbitrary new direction. Instead, the activity reset near the starting point of the original trajectory and evolved along an approximately parallel trajectory during the second (and third) sequence run (Fig. 3J). Activity was more similar between sequence runs for matched trial types (same position in sequence) than between mismatched trial types relative to shuffled controls (Fig. 3J-K, fig. S7E-F). Taken together, these results are consistent with the idea that LEC implements a flexible, multiscale representation of the timing of events.

## Slow single-cell dynamics underlying drift

We next set out to determine the mechanism underlying drift in LEC population activity. We previously found that a subset of LEC neurons exhibited slow changes in firing rate over the course of minutes (*13*). These neurons showed gradual increases or decreases in rate with a variety of time constants (i.e., “ramping” neurons), which could conceivably be driving population drift. Removing these neurons from the population, however, had no significant impact on the ability to decode time from the population, suggesting that other neurons played a more prominent role in drift.

With access to much larger populations of neurons, here we asked which neurons contributed most strongly to drift within individual foraging sessions. To do so, we calculated for each neuron the fano factor (i.e., variance over mean) of the neurońs smoothed firing rate (Gaussian width = 30 sec). This measure quantifies the variability in firing rate over a timescale of minutes, similar to the observed population drift. Activity traces for LEC neurons with high variability took a variety of forms, including ramping, multi-peaked activity, and transitions between sustained periods of (in)activity (Fig. 4A, top). The remaining neurons exhibited substantial rate changes at fast timescales yet maintained a stable mean firing rate over minutes (i.e., low variability; Fig. 4A, bottom). LEC neurons had significantly higher variability than neurons in MEC and CA1 (Fig. 4B), mirroring the differences in population drift shown above. Moreover, slow dynamics were not restricted to a defined subset of LEC neurons, but rather were broadly distributed throughout the population.

**Fig. 4.**
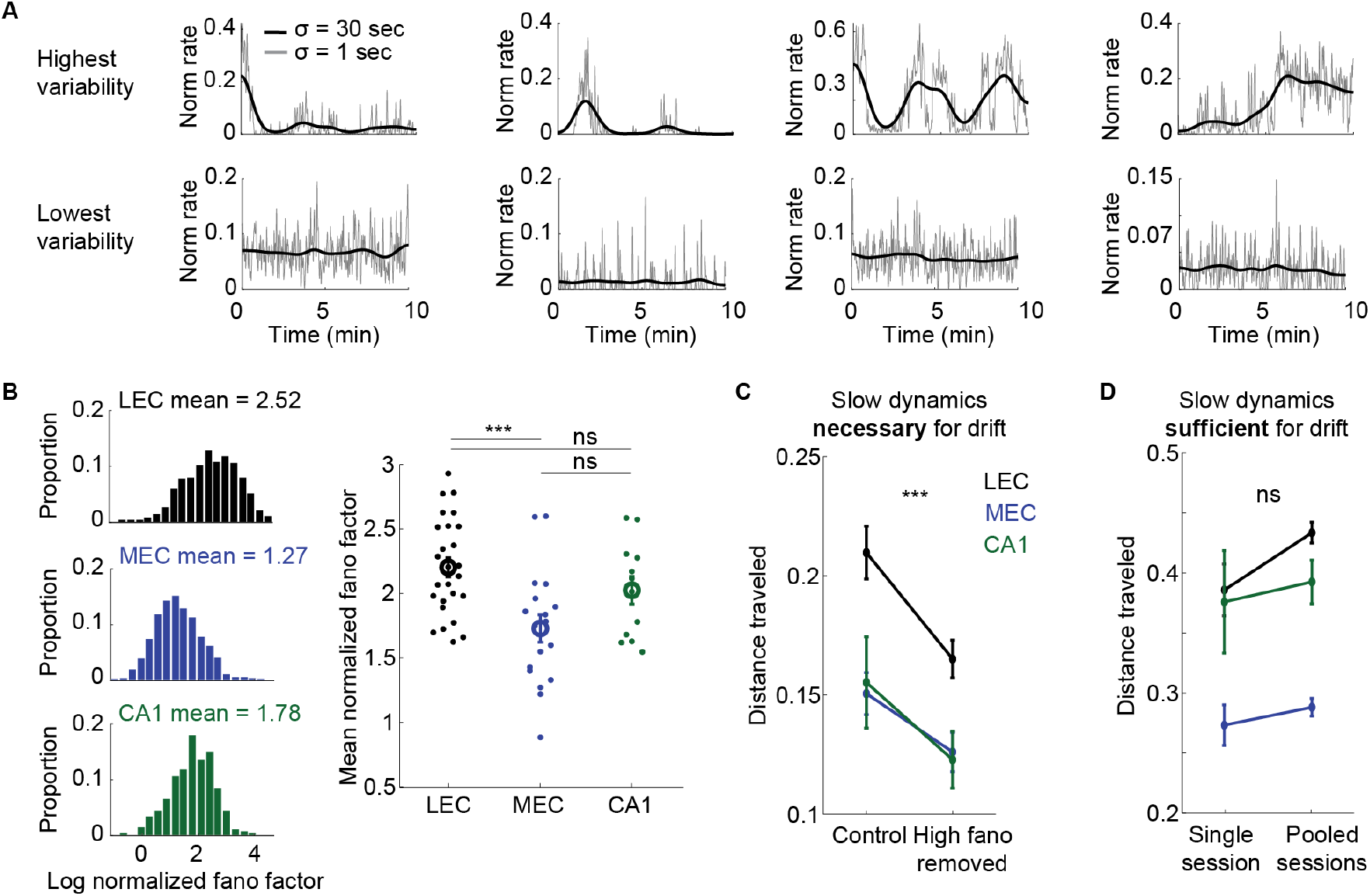
Slow dynamics in individual neurons underlying population drift. (**A**) Activity traces for the four LEC neurons with highest (top) and lowest (bottom) levels of firing rate variability over a scale of minutes in the foraging task. Firing rates smoothed with 30-sec (black) or 1-sec (gray) Gaussian. (**B**) Minute-scale variability for each neuron in an example foraging session (left) and mean variability across neurons within each session (right) for each brain area. Variability over minutes quantified as log fano factor normalized to a homogeneous Poisson neuron such that 0 is same as Poisson. LEC vs MEC: t(42) = 3.79, p = 4.80e^-4^, LEC vs CA1: t(36) = 1.36, p = 0.18, MEC vs CA1: t(28) = -1.89, p = 0.07; two-sample t-test; n = 26, 18, and 12 sessions for LEC, MEC, and CA1, respectively. (**C**) Distance traveled after subsampling from all cells within a session (Control) or from a population that had the top 25% most variable cells removed (High fano removed). Main effect between Control and High fano removed: F(1, 53) = 80.2, p = 3.47e^-12^; repeated measures ANOVA; n = 26, 18, and 12 sessions for LEC, MEC, and CA1, respectively. (**D**) Distance traveled after subsampling the top 25% most variable cells within a session (Single session) or from those same neurons pooled across all sessions (Pooled sessions). Drift was preserved after pooling which confirms that unique experiences and specific cell-to-cell correlation structure is not required for drift. Main effect between Single and Pooled: F(1, 108) = 3.8, p = 0.05; two-way ANOVA; n = 26, 18, and 12 sessions for LEC, MEC, and CA1, respectively. (**B-D**) Data represented as mean +/- SEM. ***p < 0.001, ns = not significant.

To test whether these slow dynamics in individual cells were necessary for population drift, we removed the top 25% most variable neurons (high fano removed) or removed a size-matched random sample of neurons (control) and then recomputed the total distance traveled of the neural trajectory. Removing the most variable neurons led to a significant decrease in drift (Fig. 4C), indicating that slow dynamics in single cells are required for drift in the population. To test whether these slow dynamics were sufficient for population drift, we subsampled either the top 25% most variable neurons within a single session, or a size-matched sample of those same neurons pooled across all sessions, and recomputed the total distance traveled. This pooling effectively destroys the true correlation structure of the network and averages away sensory inputs that are unique to each experience, yet the amount of drift was not reduced (Fig. 4D; fig. S8A-B). To further demonstrate that slow dynamics are necessary and sufficient for drift, we simulated networks of independent neurons that each exhibited slow dynamics and found that such networks also drift at the population level (fig. S8C-E). Taken together, the findings suggest that drift is driven by slow dynamics in neurons broadly distributed in the population.

## Synchronous ensemble responses underlying shifts

Finally, we searched for the mechanism underlying shifts at event boundaries. Because our event boundaries across different tasks took various forms at multiple timescales, we wondered whether there was a general mechanism that could elicit discrete shifts in population activity for the boundary between any two events. The discreteness implies fast dynamics in individual neurons, perhaps via boundary-induced increases in firing rate (*13, 21*). It remains unknown, however, the fraction of the population that responds at the boundary, the form of the responses, and whether the responses are stable or unique across repeated encounters with the same boundary.

To answer these questions, we characterized the responses of individual neurons on a trial-by-trial basis for each trial-based task described above. In the figure-eight task, 27% of neurons exhibited large increases or decreases in firing rate coincident with the shift in population activity as the rat approached the reward (Fig. 5A, left). In terms of both the fraction of responding neurons and the magnitude of the rate changes, single neuron responses were largest at the time of the shift compared to all other timepoints (Fig. 5A, right). Similar results were obtained in the odor sequence task where small fractions of positively or negatively modulated neurons responded at the time of the population shift as the rat approached the reward (Fig. 5B). In the novel objects task, there was a larger fraction of neurons that were positively modulated and a smaller fraction of neurons that were negatively modulated when the rats made first contact with each novel object (Fig. 5C). Lastly, we calculated the trial-to-trial variability in firing rate for boundary-modulated neurons to assess whether they could potentially barcode individual events by generating unique activity patterns at the event boundaries. We restricted this analysis to the figure-eight task where each event boundary within a session is identical (i.e., same reward every lap). Boundary-modulated responses were significantly more variable across trials compared to responses in subsets of neurons that preferred times other than the shift (Fig. 5D-E). This suggests that while boundary-modulated neurons have similar activity on average across similar events (Fig. 5A-C), they also display enough trial-to-trial variability to assign a unique barcode to each individual event (Fig. 5D). Indeed, we could successfully decode individual events (trials) using a linear decoder on held out data (Fig. 5F). Taken together, these observations point to a general mechanism for the discretization of experience whereby groups of neurons exhibit synchronous responses at event boundaries and thereby drive shifts in population activity.

**Fig. 5.**
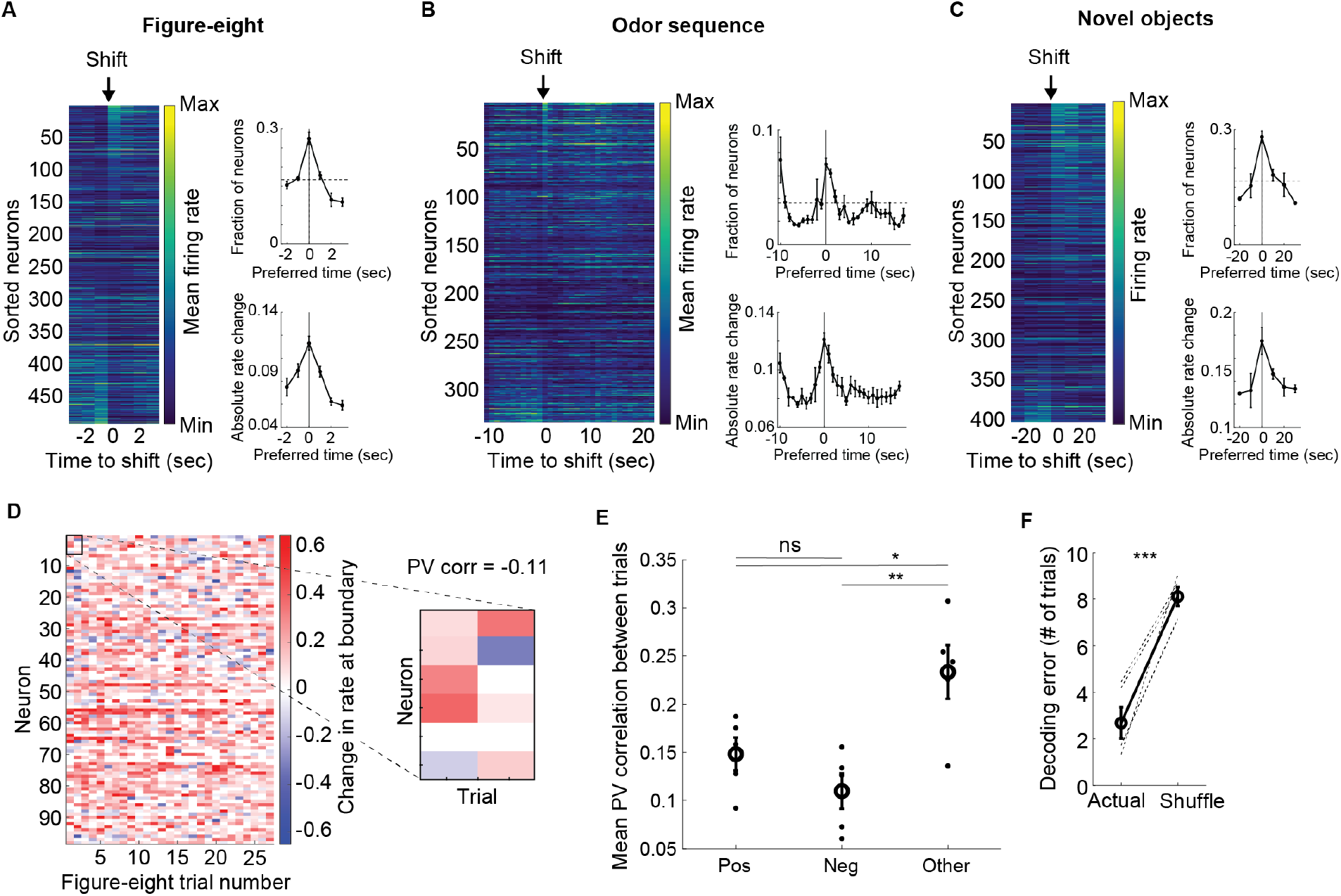
Fast dynamics in individual neurons underlying shifts at event boundaries. (**A**) Example heat map (left) of trial-averaged firing rate for all LEC neurons relative to reward in figure-eight task. Sorting neurons by rate change at the time of population shift reveals both positively and negatively modulated neurons. Across all sessions, more neurons had their largest rate change at the time of the shift compared to all other trial times (top right). Horizontal line indicates chance level of preferring each time bin. The absolute rate change was also highest for neurons that changed most at the time of the shift compared to all other times (bottom right). (**B**) Example heat map (left) and summary data (right) showing neurons modulated at the reward-related population shift in the odor sequence task. The initial peak on the left corresponds to the other population shift when the treadmill turns on. Conventions as in (A). (**C**) Example heat map (left) and summary data (right) showing neurons modulated at the contact-related shift in the novel objects task. Conventions as in (A). (**A-C**) Modulated neurons defined as those with their largest absolute change in rate (relative to previous time bin) at the shift time (vertical lines). Chance level fractions (horizontal lines) were 17%, 4%, and 17% for figure-eight, odor sequence, and novel objects tasks, respectively, based on the number of time bins in each task. (**D**) Example data from figure-eight session showing that even after selecting for neurons with trial-averaged increases in firing rate at the event boundary, this subset showed substantial trial-to-trial variability. Note the blue areas showing individual trials when these positively modulated neurons actually showed negative responses. The inset to the right highlights six example neurons for the first two trials with a population vector (PV) correlation of -0.11, demonstrating that LEC activity creates unique barcodes for each trial. (**E**) Mean PV correlation between responses at event boundaries during figure-eight trials for positively modulated neurons (Pos), negatively modulated neurons (Neg), and randomly chosen subsets (n = 25) of neurons that preferred times other than the shift (Other). Pos vs Neg: t(8) = 1.53, p = 0.17, Pos vs Other: t(8) = -2.60, p = 0.03, Neg vs Other: t(8) = -3.70, p = 6.01e^-3^, two-sample t-test; n = 5 sessions. (**F**) Unique barcodes for each event enabled accurate decoding of trial number using a linear classifier trained on held out data. Decoding error is measured as the difference between actual and predicted trial number. Shuffle obtained by shuffling epoch labels. Actual vs shuffle: t(4) = -8.64, p = 9.89e^-4^, paired t-test; n = 5 sessions. (**A-C, E-F**) Data represented as mean +/- SEM. ***p < 0.001, **p < 0.01, *p < 0.05, ns = not significant.

## Discussion

Leveraging the power of high-density, multi-area unit recordings in freely behaving rats, we have shown how event structure impacts neural population dynamics in LEC. When experimental conditions were stable, neural population activity in LEC, but not MEC or CA1, drifted progressively along a non-periodic one-dimensional manifold in the population state space, regardless of the animal’s behavior or sleep-wake state. By doing so, the activity functioned as a population clock (*31*), allowing downstream neurons to read out the passage of time during experience (*7*). This continuous drift was interrupted, however, by abrupt shifts at event boundaries, including salient changes in the experimental context such as encountering a reward or a novel object. These shifts introduced non-linearities in the neural trajectory, segmenting the stream of experience into discrete events which could later be recalled as individual units (*7, 24*). The combination of drifting and shifting dynamics within a single neural population during the stream of experience may be passed on to hippocampal memory circuits and underlie temporal memory judgements as well as distortions in temporal memory imposed by event boundaries (*17*). The data are consistent with observations in fMRI studies of human hippocampus where sharp onset and offset responses at event boundaries (*16, 21–23*), similar to the shifts described in neural spike data here, have been identified as a potential source of bias in future judgments of temporal order (*21*). Our results show that LEC has a mechanism for encoding event boundaries as discontinuities in the progressive drift of neural population activity in LEC. Discontinuities were associated with bursts of activity in ensembles of LEC cells. How such discontinuity-generating bursts arise remains to be determined, yet it is known that LEC activity is strongly modulated by novelty-related prediction error signals, such as those emerging from neuromodulatory systems in the brain stem (*32–35*). Prediction error signals to LEC could shift trajectories in state space, which in turn may account for distortions in memory for temporal duration when experiences span event boundaries (*36–38*).

Our findings yield insight into the geometry of drift in neural state space. The population activity did not simply drift uniformly across the session. During tasks with repetitive temporal structure, activity also traveled (at a faster timescale) in directions orthogonal to the drift, enabling the system to multiplex temporal information about the task with the slower changes reflecting session time. By traveling in independent directions for each behaviorally relevant timescale, the neural code in LEC is thus inherently multiscale, with fast timescales nested inside slow timescales. Multiscale temporal coding has been studied extensively at the level of neural oscillations (*39, 40*), but few studies have explored longer timescales of seconds to minutes. The present findings show that activity can progress along multiple timescales also under non-periodic conditions. Individual neurons were not obviously locked to a single preferred timescale but instead flexibly multiplexed different timescales of experience, allowing LEC and readout neurons to continuously capture the temporal statistics of the task. This hierarchical representation in LEC could facilitate the ability to deconstruct events (*17*) into sub-events at timescales ranging from seconds to minutes or more and thus enable the flexible examination of memories at a level appropriate for the current goals.

Lastly, we identified candidate mechanisms underlying drift and shifts that provide insight into how a multiscale temporal code of experience may be generated. Drift was associated with slow, minute-scale variability in the firing rates of individual neurons. While a subset of these neurons displayed gradual ramping with different time constants (*13, 41*), most neurons had a variety of other forms of slow dynamics. We confirmed in network simulations that the richness of these firing rate dynamics in LEC, i.e., the variability over time and between neurons in this area, is the main driver of the slow drift in neural population space. In contrast to drift, shifts in state space were associated with synchronous responses of groups of neurons at event boundaries. These synchronous responses may be elicited by external inputs targeting subsets of LEC neurons, which are known to encode stimuli of a wide variety of sensory modalities, reflecting LEC’s position as a node of convergence for inputs from widespread cortical and subcortical regions (*42, 43*). They might also be induced by neuromodulatory inputs from the brainstem associated with novelty and prediction error (*32–35*). Boundary responses showed some degree of generalization across similar events, but the combination of activated cells varied sufficiently between boundaries such that LEC populations, much like those in CA1 (*44, 45*), collectively barcoded the individual events. When read out by neural circuits downstream in the hippocampus, unique barcodes from LEC at event boundaries may be stored and retained as individual, orthogonalized episodic memories (*44–47*). These memories may then form the basis for estimates of duration and temporal order during recall of experiences (*7*).

## Acknowledgments

We thank A.M. Amundsgård, S. Ball, K. Haugen, E.H. Holmberg, K.J. Jenssen, E. Kråkvik, H. Waade for technical assistance, the veterinary staff for animal care, R. Gardner and V.A. Normand for initial training with Neuropixels, and B.A. Dunn, J.A. Gallego, R. Gardner, I. Polti, J. Sugar, A.Z. Vollan, and T. Waaga for discussion.

## Funding

ERC Synergy Grant 951319 (EIM)

Centre of Neural Computation 223262 (EIM, MBM)

Centre for Algorithms in the Cortex 332640 (EIM, MBM)

Kavli Foundation

Ministry of Science and Education, Norway

## Author contributions

Conceptualization: EIM, BRK

Methodology: BRK, CML, EIM, MBM

Software: BRK

Formal analysis: BRK

Investigation: BRK, CML

Data curation: BRK

Writing – original draft: BRK

Writing – review & editing: BRK, EIM, MBM, CML

Visualization: BRK, CML

Supervision: EIM, MBM

Project administration: EIM, MBM

Funding acquisition: EIM, MBM

## Data and materials availability

The datasets generated during the current study and the code for reproducing the analyses will be available before publication.

### List of Supplementary Materials

Materials and Methods

Table S1

Figs. S1 to S8

## Materials and Methods

### Subjects

The data were collected from seven adult male Long Evans rats weighing approximately 400– 500 g at time of implantation. The rats were group-housed with one to eight of their male littermates before surgery and were housed alone in a large two-story enriched metal cage (95 x 63 x 61 cm) thereafter. They were kept on a 12-h light–12-h dark schedule, with strict control of humidity and temperature. All experiments were approved by the Norwegian Food Safety Authority (FOTS ID 18011) and performed in accordance with the Norwegian Animal Welfare Act and the European Convention for the Protection of Vertebrate Animals used for Experimental and Other Scientific Purposes.

### Surgery and Electrode Implantation

Rats were implanted with Neuropixels 2.0 silicon probes targeting LEC, MEC, and/or CA1. One rat (27284) had a Neuropixels 1.0 single-shank probe implanted in medial entorhinal cortex. LEC probes were implanted 6.23-7.00 mm posterior to bregma, 3.70-4.05 mm lateral of the midline, at an angle of 20 deg in the coronal plane with the tip of the probe pointing laterally. Probes were lowered until one or more shanks met resistance at the ventral surface, and were then retracted 100 microns, reaching a final depth 7.84-9.11 mm below the pial surface. MEC probes were implanted 100 microns anterior to the transverse sinus, 4.60 mm lateral of the midline, at an angle of 26 deg in the sagittal plane with the tip of the probe pointing anteriorly. Probes were lowered 5.50 mm below the pial surface. CA1 probes were implanted 3.80 mm posterior to bregma and 3.20 mm lateral of the midline. Probes were lowered 6.00-6.19 mm below the pial surface. Table S1 reports the exact implant coordinates for each probe in each rat. The implant was secured with dental cement. A small stainless-steel screw was attached to the skull above the cerebellum and connected to the probe ground and external reference pads with insulated silver wire. See (*25*) for further details of probe implantation. After surgery, rats were left to recover until resuming normal locomotor behavior, a minimum of 2 hours. Postoperative analgesics (meloxicam and buprenorphine) were administered during recovery.

### Recording Procedures

The details of the Neuropixels hardware system and the procedures for freely moving recordings have been described previously (*25, 48*). In brief, electrophysiological signals were amplified with a gain of 80, filtered 0.005-10 kHz, and digitized at 30 kHz by the probe’s on-board circuitry. The digitized signals were multiplexed by an implant-mounted headstage circuit board and were transmitted along a lightweight 5-m tether cable, made using twisted pair wiring. SpikeGLX software (https://billkarsh.github.io/SpikeGLX/) was used for data acquisition and configuring the probes. Three-dimensional motion capture (OptiTrack Flex 13 cameras and Motive recording software) was used to track the rat’s head position and orientation by attaching a set of five retroreflective markers to the implant during recordings. The 3D marker positions were projected onto the horizontal plane to yield the rat’s 2D position and head direction. An Arduino microcontroller was used to generate digital pulses, which were sent to the Neuropixels acquisition system (via direct TTL input) and the OptiTrack system (via infra-red LEDs) to permit precise temporal alignment of the recorded data streams.

### Behavioral Procedures

Data were obtained from several recording sessions performed within the first week after recovery from surgery. Recordings were performed while the rats engaged in four behavioral paradigms (or sleep sessions) using multiple mazes/arenas and rooms. Many distal visual and auditory cues were available to the rat. During pre-surgical training and habituation, several of the rats were food-restricted through intermittent fasting during which food was available ad-libitum for four hours between 12:00 and 17:30. During that training phase, behavioral procedures were done from 8:00 when the animals were maximally food motivated. Food restriction ceased a minimum of 24 hr prior to surgery.

### Foraging Task

Rats foraged for randomly scattered food crumbs (corn puffs and vanilla cookies) in a square open-field arena with a diameter of 1, 1.5, or 2 m. The arena had dark blue or black flooring and was enclosed by walls of height 50 cm. Large distal cues were available outside of the arena near the room walls. The arena was dimly lit by one or two lamps along the room wall. At the time of surgery, four rats (27284, 27285, 27963, and 28003) were familiar with the environment and task (minimum four x 10 min sessions). Three rats (26863, 26965, and 26966) were completely naïve to the arena, room, and task at the time of their first recording session. Recording sessions lasted between 12 and 142 min.

### Figure-eight Task

Two rats (26965 and 26966) were trained to run laps around a figure-eight maze, receiving one reward per lap. The maze was made of wood with vinyl flooring and plastic lips (2 cm high) and was elevated 80 cm above the ground by metal table legs. After being placed at the base of the maze, rats ran down a 50 cm long (12 cm wide) central stem to the top of the maze which was a 50 x 50 cm square with a small reward port (polyurethane tubing leading to 15 ml conical tube cap) at the far end. After drinking a sweetened chocolate milk reward (2.5% sucrose in Oatly chocolate milk), the rat could run back along either return arm (12 cm wide) to reach the base of the central stem again. The maze was open to the room with many available distal cues. The room was dimly lit by two small lamps on the left room wall. Animals were prevented from running in the wrong direction using a tall plastic barrier during training. During training and testing, a large plastic door was also used at the top of the central stem to prevent backtracking. The door opened as the rat came down a return arm and closed again after the rat retrieved a reward. Animals were considered trained when performing approximately 20 trials per session for multiple days and were implanted shortly thereafter.

### Odor Sequence Task

One rat (27285) was trained to run laps around the same figure-eight maze described above, with a few small modifications. A milk port was added at the base of the central stem. The central stem itself was a treadmill with a large front door to prevent the rat from leaving until the treadmill turned off. At the top of the maze, the rat was presented with an odorized cup of sand containing a buried chocolate cookie crumb reward (ChocoLoops). Odors were 1 of 10 common household spices, thoroughly mixed in sand with the following concentrations: A = parsley, 1%; B = cumin, 0.5%; C = paprika, 1%; D = thyme, 1%; E = cardamom, 0.8%; L = clove, 0.5%; M = tarragon, 1%; N = cinnamon, 0.8%; Ø = dill, 1%; P = coffee, 1%. A custom GUI written in MATLAB was used to control the treadmill, door, and milk delivery. Each trial began when the rat reached the end of the treadmill, which triggered the treadmill to turn on at 30 cm/sec for 10 sec. After 10 sec, the treadmill turned off and the large front door opened so the rat could run to the sand cup to dig for a reward. The rat then ran via either return arm to receive a sweetened chocolate milk reward (2.5% sucrose in Oatly chocolate milk) at the base of the central stem, before entering the treadmill again to initiate another trial. On each trial the odor in the sand was different, creating a sequence of five odors A through E across five trials. Sand and odors from the previous trial were removed with a handheld vacuum during the 10 sec treadmill run, after which the sand cup for the next trial was put in position. This ensured that the rat could not smell the upcoming odor or chocolate treat until after leaving the treadmill. These five trials comprised a run, and the rat ran three runs with 5 min rest in a flowerpot between each run, and also before and after the runs: rest, RUN1, rest, RUN2, rest, RUN3, rest. The rat ran this sequence (SEQ1) in the morning, and after a 2 hr delay in the home cage, returned to run a second sequence (SEQ2) in the afternoon. The only difference between morning and afternoon is that SEQ2 contained five different odors L through P. Data from the two different sequences were treated equivalently for analysis purposes. Shaping to dig and run laps took several days. Training on the full task with both sequences occurred over several days and surgery was conducted after training day 3. Recordings lasted approximately 30 min for each sequence.

### Novel Objects Task

Two rats (27963 and 28003) foraged for randomly scattered food crumbs (corn puffs and vanilla cookies) in a square open-field arena, exactly as in the foraging task described above. After 7.5 min of foraging in the empty arena, a novel object was placed at a pseudorandom location on the floor of the arena. Objects had footprints of approximately 10 x 10 cm and were approximately 15 cm tall. They consisted of common laboratory or household items (e.g., beaker, flask, spray bottle, candlestick). 7.5 min later that object was removed and another novel object was placed at different pseudorandom location. This was repeated once more 7.5 min later resulting in the following sequence: Empty, OBJ1, OBJ2, OBJ3. Recordings therefore lasted 30 min. At the time of surgery, rats were familiar with the environment and foraging task (minimum four x 10 min sessions) but had never experienced objects in the arena before. Both rats went through the OBJ experiment several times in different recording sessions.

### Natural Sleep

Two rats (27284 and 27285) were recorded during natural sleep by placing them in dedicated sleep box made of black acrylic (30 × 30-cm floor, 80 cm height). The floor contained a shallow flowerpot lined with several towels to make a nest and rats were habituated to the box over a minimum of four sessions before implantation. The box walls passed infrared light to enable tracking through the walls. Full room lights were on and pink noise was played through the computer speakers at approximately 60 dB to mask background sounds. Sleep sessions were conducted at the end of the light phase (7:00-8:00) and lasted between 45 and 140 min.

### Spike Sorting and Unit Selection

Spike sorting was performed with KiloSort 2.5 with customizations as previously described (*48*). Units were discarded if more than 2% of their interspike interval distribution consisted of intervals less than 2 ms. In addition, units were excluded if they had a mean spike rate of less than 0.05 Hz or greater than 40 Hz (calculated across the full recording duration), or if they were recorded on sites outside the region of interest.

### Preprocessing and Temporal Binning

Data was not filtered for running speed. Spikes were binned using 0.5-10 sec time bins, depending on the timescale of interest for each task, and tracking data was resampled at the same time intervals to align it with the spike data. Spike count vectors for each neuron were ‘soft’ normalized (*49*) to reduce the impact of strong responses by dividing the counts by the range of counts + 5, where 5 is the normalization factor. Spike time matrices for each region consisted of all units that met the selection criteria above.

Neural populations were not pooled across recording sessions. By restricting the analysis to populations of simultaneously recorded neurons, we avoided potential spurious results caused by mixing recording sessions of neural activity in different functional modes. The one exception to this rule is the analysis presented in Fig. 4D which shows that pooling neurons across recording sessions does not eliminate drift at the (pseudo)population level.

### Sleep Stage Classification

Sleep stages were identified as described previously (*48*). Periods of sustained immobility (lasting > 120 sec, locomotion speed < 1 cm/sec, head angular speed < 6 deg/sec) were classified into SWS and REM based on the theta/delta ratio of MEC population activity. Periods when the theta/delta ratio remained above 5.0 for at least 20 sec were classified as REM, whereas periods when the theta/delta ratio remained below 2.0 for at least 20 sec were classified as SWS. Only REM periods were further analyzed.

### Dimensionality Reduction

Dimensionality reduction was used mainly for visualizing neural trajectories. It was additionally used for quantifying tangling of trajectories and decoding temporal epochs. Distances in state space were always calculated in the full dimensional space (see below) to avoid potential distortions in lower dimensional embeddings.

Principal component analysis (PCA) was run on the soft normalized spike × time matrices for each region using the sci-kit learn function *PCA*. Linear discriminant analysis (LDA) was used to find dimensions capturing change over time using the sci-kit learn function *LinearDiscriminantAnalysis*. Principal components explaining 50% of the variance were used as input to LDA and class labels were defined as temporal epochs, described above for individual tasks.

### Distance Traveled and Change in Trajectory Speed

Distance traveled was defined as the cosine distance (using the SciPy function *pdist*) between population vectors in the full dimensional space of N neurons. For comparisons between temporal epochs consisting of multiple time bins, distance was calculated as the mean pairwise distance between the epochs.

Change in trajectory speed (acceleration/deceleration) was defined as the second derivative of distances between population vectors for neighboring time bins. Agnostic search for event boundaries was done by calculating the instantaneous change in trajectory speed throughout the whole recording session. Putative times for discrete shifts in state space were defined as those when the population firing rate exceeded the 90^th^ percentile and the increase in population firing rate from the previous time bin also exceeded the 90^th^ percentile (different choices of threshold yielded similar results). Change in trajectory speed at those onsets of synchronous activity were compared to all other times in the recording session.

### Tangling of Neural Trajectories

Tangling of neural trajectories was calculated in the 2D space of the top 2 linear discriminants following PCA/LDA as described above. Tangling was defined as in (*50*):

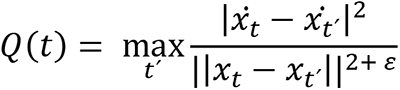

where *x*_*t*_ is the population vector at time *t*, 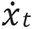 is the temporal derivative of the neural state, ||⋅|| is the Euclidean norm, and *ε* is a small constant that prevents division by zero.

### Decoding of Temporal Epochs

Decoding of temporal epochs was in the full space of linear discriminants following PCA/LDA as described above. Data was split into five cross validation folds using the sci-kit learn function *KFold*. For decoding time within a session, time bins were shuffled before splitting into folds so that training data consisted of time bins from several temporal epochs. For decoding time within a trial or trial type, data was split into five folds based on trials such that entire trials were held out of the training data. Temporal epochs were predicted for each time bin using the sci-kit learn function *cross_val_predict*. Decoding accuracy was defined as the percentage of correctly predicted epochs, averaged over the five folds, using the sci-kit learn function *accuracy_score*. Decoding error was defined as the mean difference in time between predicted and actual epochs.

### Comparison of Drift during REM Sleep and Wake

Drift during REM sleep was compared to drift during wake by creating size-matched spike count matrices. Two rats (27284 and 27285) with probes in both LEC and MEC (used for detecting REM periods) with sleep sessions containing several extended REM periods were included in this analysis. Because typical REM periods lasted only a few minutes, periods of 2.5 min were used to assess drift in both REM sleep and awake foraging. REM periods were truncated by keeping the first 2.5 min of candidate REM periods. Foraging periods were truncated by dividing the 10-min foraging session into four equal parts. Data for REM and foraging were analyzed using 5-sec temporal bins and 30-sec epochs. Distance traveled during REM periods was defined (as above) as the mean pairwise cosine distance between first and last temporal epochs. Distance traveled during foraging was calculated in the exact same manner, but distances were further averaged across the four equal parts of a continuous foraging session to avoid ‘double dipping’ from the same data.

### Definition of Trial Data from Continuous Behavior

Trials in the figure-eight task were defined post-hoc from continuous lap running behavior. Trials were aligned based on the x-y position of the rats (from head-mounted markers) just before stopping to consume the reward. This point was calculated by finding the mode of the distribution of all y-position values throughout the session when the rat was within a defined x-position range capturing the central stem. It was confirmed by manual inspection that this corresponded to the location of reward consumption. The trial alignment point was then defined as 3 cm below that location to ensure exclusion of reward consumption itself in some analyses. For analysis of shifts in state space (Fig. 2), trials were defined from 3 sec before the alignment point to 4 sec after (including reward). For analysis of within-trial time (Fig. 3), trials were defined as the 6-sec periods leading up to the alignment point (excluding reward).

Trials in the odor sequence task were defined based on logged timestamps of the treadmill turning on. Each trial started when the treadmill turned on and ended either when the treadmill turned on for the next trial, or when it was the final lap of the sequence run, after consuming the reward at the top of the maze (thus the final laps were shorter than the rest). For comparing activity across trials, trials were truncated to the fastest full lap. For observing the full dynamics in Fig. 3J, all times during sequence runs and intertrial intervals were included.

### Neural Trajectory Alignment

Neural trajectory alignment was used to assess whether matched trial data was closer together in state space compared to mismatched trial data. Alignment was defined as:

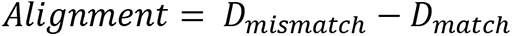

where *D_mismatch_* is the mean pairwise cosine distance between time bins with mismatched temporal epochs and *D_match_* is the same but for time bins with matched temporal epochs. Alignment values were rather small because mismatched distances included comparisons of neighboring temporal epochs where distances are expected to be small.

### Angular Offset between Coding Dimensions

The angular offset between coding dimensions (e.g., session or trial time) was defined by running PCA/LDA, as described above, for each coding dimension (CD) separately. The PC with the largest contribution to the top LD was identified and its PC loadings were extracted. The same procedure was done for the other CD. Angular offset was defined as:

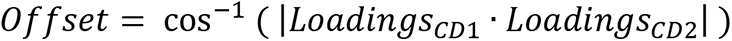

The probability that any pair of vectors is orthogonal increases in higher dimensions. To avoid spurious claims of orthogonality, a leave-one-out resampling procedure was used to quantify the stability of each CD over time. For session time, loadings were re-calculated after leaving out one temporal epoch. This was repeated for each temporal epoch. Within CD stability was defined as the mean angular offset between each of the resampled vectors and the original vector. For trial time, loadings were re-calculated after leaving out one trial. This was repeated for each trial. Within CD stability was again defined as the mean angular offset between each of these resampled vectors and the original vector.

The angular offset approach was well-suited to identifying the best CD vector for each timescale yet it required two separate sources of input data (full session data with 10-sec bins versus trial-based data with 1-sec bins). A complementary approach allowed us to observe the orthogonal CDs in a single common subspace. PCA was run on the trial-based data with 1-sec bins. Each of the top PCs were then examined to check for strong correlations with either trial time or session time. When such correlations were present, we could visualize both coding dimensions in a single 2D subspace defined by those PCs, which are orthogonal by definition.

### Multiplexing of Coding Dimensions

Multiplexing of coding dimensions (e.g., session or trial time) was defined by running PCA/LDA, as described above, for each coding dimension (CD) separately. The PC with the largest contribution to the top LD was identified and its absolute PC loadings were extracted. The same procedure was done for the other CD. These loadings were plotted against each other for visualization and neurons exceeding the 75% percentile of both distributions of loadings were considered as potential multiplexing neurons (i.e., displaying mixed selectivity) (fig. S6H). Neurons at the extreme ends of both distributions of loadings were further visualized as a proof of principle that multiplexing of these timescales is possible (fig. S6I).

### Minute-scale Variability in Neural Firing Rates

Individual neuron spike trains during 10-min foraging sessions were binned in 0.5 sec bins and then smoothed with a Gaussian of width σ = 30 sec. The fano factor of this smoothed firing rate vector was defined as:

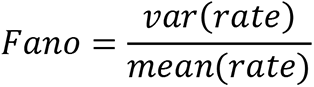

To compare these values to a known reference, simulated spike trains were sampled from a homogeneous Poisson process and fano factors were calculated on these simulated Poisson neurons in the same manner. Log normalized fano factor values reported in Fig. 4 were obtained by dividing the fano factor of each real neuron by the mean fano factor of 500 simulated Poisson neurons, and then taking the log of this value. Based on this normalization, a value of 0 indicates the same amount of variability as observed in Poisson neurons.

### Correlation Structure

Pairwise correlations were calculated as the Pearson correlation between all pairs of smoothed firing rate vectors during individual 10-min foraging sessions (as defined in previous section).

Breaking correlation structure in fig. S8B was done by circularly shifting unsmoothed firing rate vectors for simultaneously recorded neurons relative to each other. Each neuron was shifted in time independently by a random interval between -2 min and 2 min. The first and last 2 min of the spike × time matrix was then truncated to eliminate edge effects from the shifting procedure, and distance traveled during the remaining 6 min of the foraging session was calculated as above.

### Network Simulations of Drift

Simulated spike trains (n = 500 units) were sampled from a homogeneous Poisson process, as described above. Each simulated unit was then duplicated such that one copy was smoothed with a Gaussian of width σ = 30 sec (slow) and the other copy was smoothed with a Gaussian of width σ = 1 sec (fast). Example units are shown in fig. S8C. Neural trajectories and distance traveled over 10 min were calculated for slow and fast populations separately (fig. S8D-E) using the same methods described above. This procedure was repeated for a total of 25 simulations, the results of which are individually displayed in fig. S8E.

### Event Boundary Responses in Individual Neurons

The preferred time for each neuron in trial-based tasks was defined as the time bin with the largest absolute change in trial-averaged firing rate relative to the preceding time bin. The fraction of neurons preferring each time bin was calculated as the number of neurons preferring each bin divided by the total number of simultaneously recorded neurons (i.e., calculated per session). The absolute rate change was calculated as the mean absolute change in trial-average firing relative to the preceding time bin, calculated over all time bins, for all simultaneously recorded neurons.

Barcoding of individual events was assessed by calculating the mean population vector correlation across all figure-eight trials for different subsets of simultaneously recorded neurons. Positively modulated neurons were defined as neurons with preferred times (defined in previous paragraph) at the event boundary that had a trial-averaged increase in firing rate at that time relative to the preceding time bin. Negatively modulated neurons were defined in the same manner for neurons with decreases in firing rate. Control populations were defined as random samples of 25 simultaneously recorded neurons, and mean population vector correlations were averaged over 50 random samples. Decoding of trial identity was performed as described above (Decoding Temporal Epochs) except that trial numbers were used as epochs instead of trial times. All simultaneously recorded neurons were included in the decoding analysis. Decoding error was defined as the mean difference in trial number between predicted and actual epochs.

### Histology and Recording Locations

Rats were given an overdose of sodium pentobarbital and were perfused intracardially with saline followed by 4% formaldehyde. The extracted brains were stored in formaldehyde and a cryostat was used to cut 30-μm sagittal sections, which were then Nissl-stained with cresyl violet. The probe shank traces were identified in photomicrographs, and a map of the probe shank was aligned to the histology by using the tip of the probe shank as a reference point. The recorded area of the probe was near-parallel to the cutting plane; therefore, it was deemed sufficient to perform a flat 2D alignment in a single section. The aligned shank map was then used to calculate the anatomical locations of individual recording sites (fig. S1-2).

### Data Analysis and Statistics

Data analyses were performed with custom-written scripts in Python 3.10 and MATLAB 2023a (MathWorks). Open-source Python packages used were: NumPy, SciPy, and sci-kit learn.

Statistical analysis was performed in MATLAB. Power analysis was not used to determine sample sizes. The study did not involve any experimental subject groups; therefore, random allocation and experimenter blinding did not apply and were not performed. Error is reported as standard error of the mean. Sample sizes are reported in the Results section.

---

**Table S1.** Implant coordinates. Each row contains stereotactic coordinates for a single probe. Seven rats (separated by thick black lines) each had one to three probes implanted.

| **Animal ID–**<br>**Brain area** | **AP (mm)**<br>**reference** | **ML from**<br>**midline (mm)** | **DV from pia**<br>**(mm)** | **Angle (deg)** |
| --- | --- | --- | --- | --- |
| 26863 - LEC | -7.00, bregma | +3.70 | -7.84 | 20, tips lateral |
| 26965 - LEC | -6.60, bregma | +4.05 | -8.22 | 20, tips lateral |
| 26966 - LEC | -6.60, bregma | +4.05 | -8.39 | 20, tips lateral |
| 27284 - LEC | -6.60, bregma | -4.05 | -8.57 | 20, tips lateral |
| 27284 - MEC | 0.10, transverse<br>sinus | +4.60 | -5.50 | 26, tips anterior |
| 27285 - LEC | -6.60, bregma | -4.05 | -8.69 | 20, tips lateral |
| 27285 - MEC | 0.10, transverse<br>sinus | +4.60 | -5.50 | 26, tips anterior |
| 27285 - CA1 | -3.80, bregma | +3.20 | -6.19 | none |
| 27963 - LEC | -6.60, bregma | -4.05 | -9.00 | 20, tips lateral |
| 27963 - MEC | 0.10, transverse<br>sinus | +4.60 | -5.50 | 26, tips anterior |
| 27963 - CA1 | -3.80, bregma | +3.20 | -6.19 | none |
| 28003 - LEC | -6.23, bregma | -4.05 | -9.11 | 20, tips lateral |
| 28003 - MEC | 0.10, transverse<br>sinus | +4.60 | -5.50 | 26, tips anterior |
| 28003 - CA1 | -3.80, bregma | +3.20 | -6.00 | none |

**Fig. S1.**
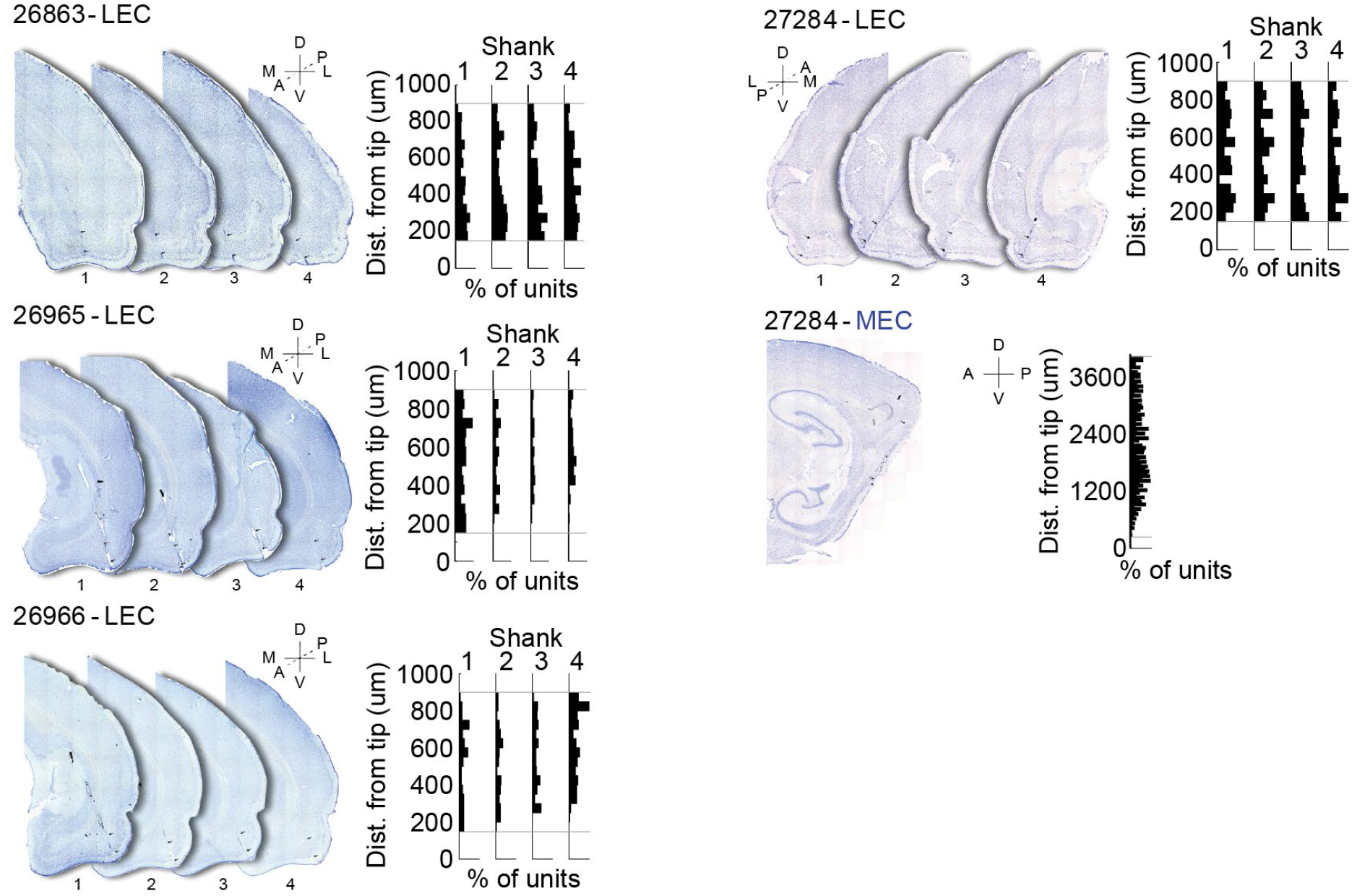
Histological confirmation of recording sites for rats with one or two probes. Coronal (LEC) and sagittal (MEC) sections stained with cresyl violet show probe placement for each brain area in each rat. Black arrows indicate the dorsal and ventral extent of recorded locations along the probe. D = dorsal, V = ventral, M = medial, L = lateral, A = anterior, P = posterior.

**Fig. S2.**
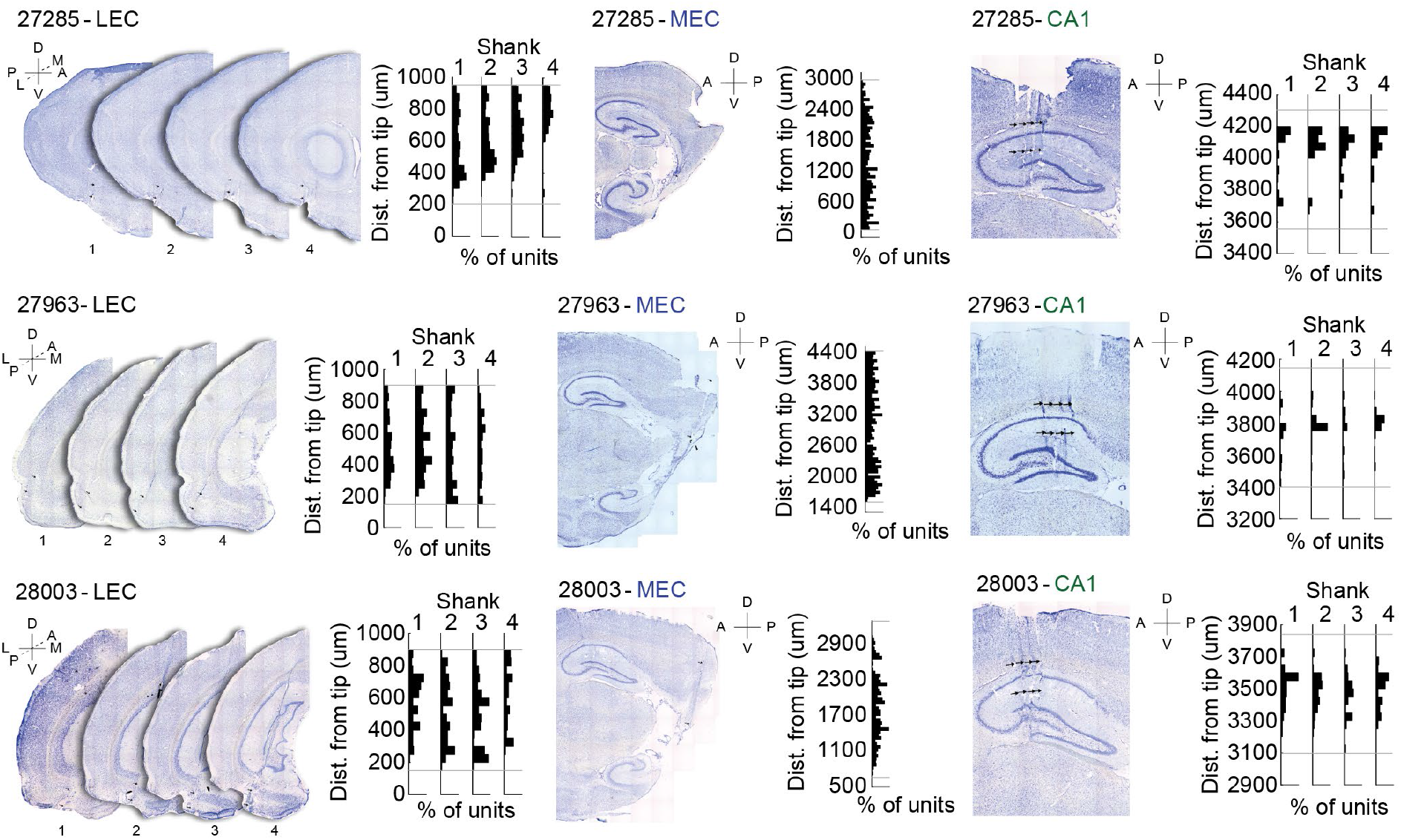
Histological confirmation of recording sites for rats with three probes. Coronal (LEC, except 27285 which is sagittal) and sagittal (MEC and CA1) sections stained with cresyl violet show probe placement for each brain area in each rat. Black arrows indicate the dorsal and ventral extent of recorded locations along the probe. D = dorsal, V = ventral, M = medial, L = lateral, A = anterior, P = posterior.

**Fig. S3.**
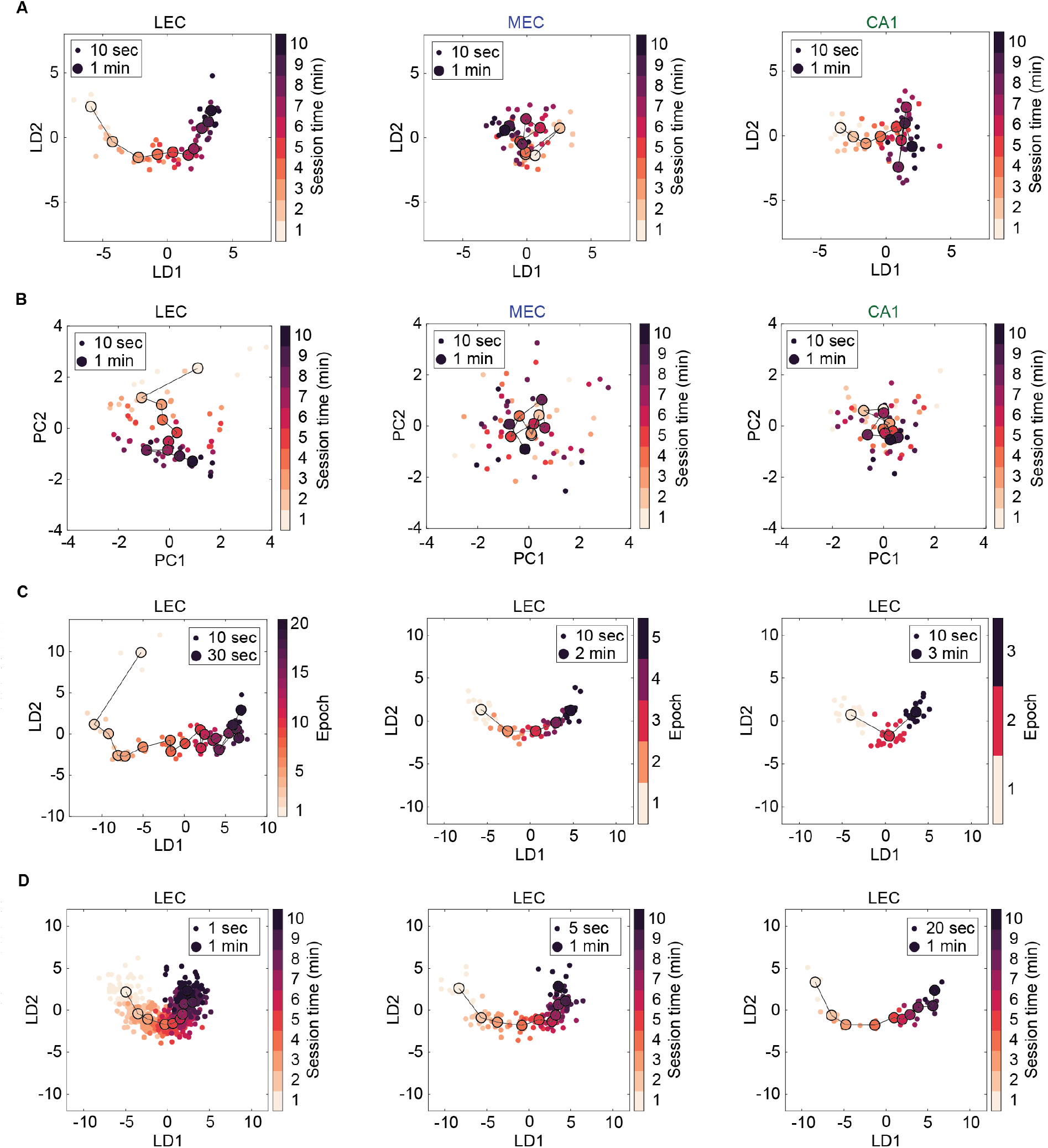
Alternative visualization of neural trajectories. (**A**) Similar trajectories obtained by applying linear discriminant analysis directly on the matrix of normalized spike counts without first applying principal component analysis. (**B**) Similar trajectories obtained by applying only principal component analysis on the matrix of normalized spike counts without subsequently applying linear discriminant analysis. (**C**) Similar trajectories in LEC obtained by using different epoch sizes as class labels for linear discriminant analysis. (**D**) Similar trajectories in LEC obtained by using different bin sizes prior to dimensionality reduction. (**A-D**) Example data (from rat 27285) and plotting conventions are the same as in Fig. 1B. Brain region indicated above each panel.

**Fig. S4.**
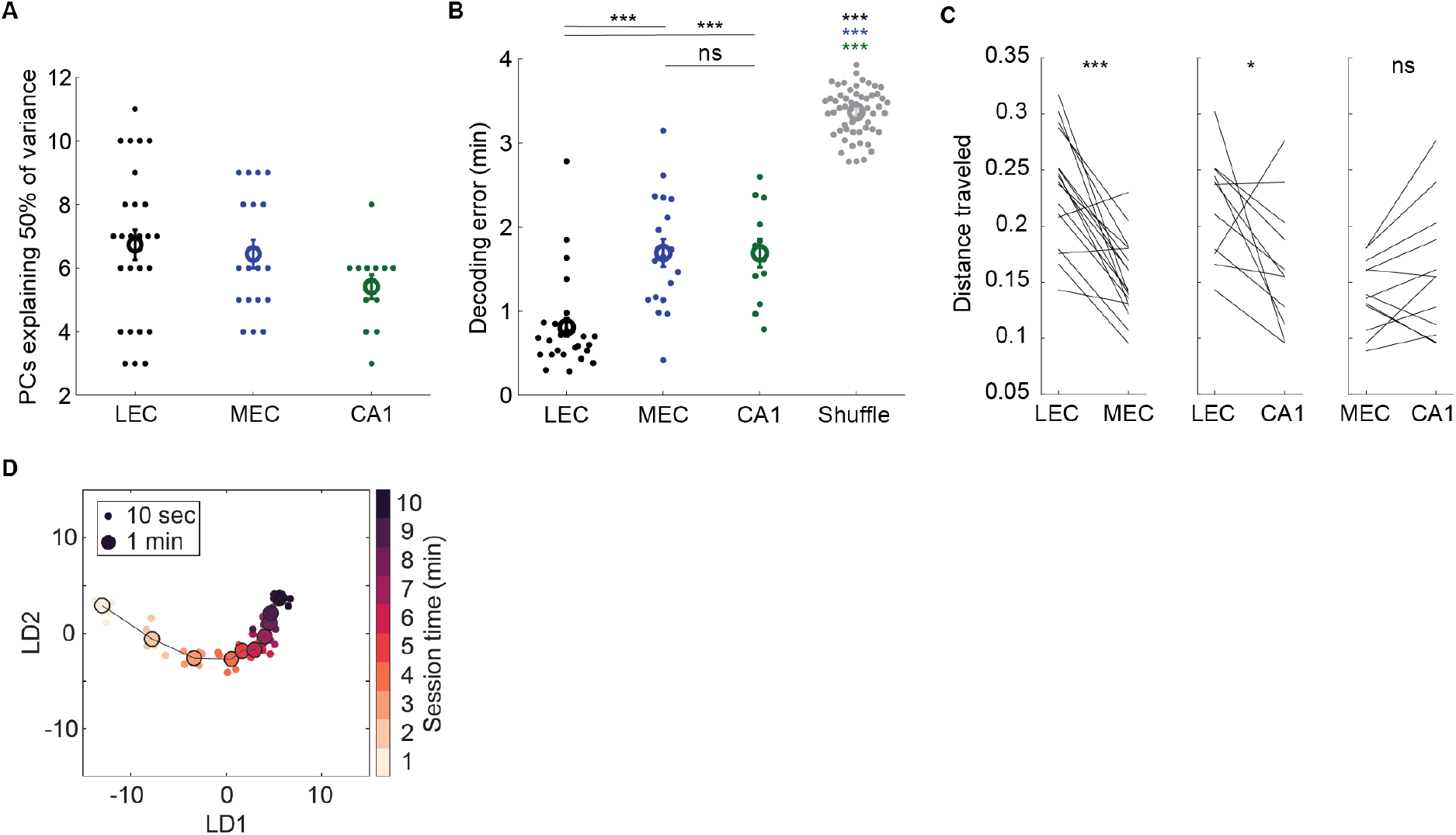
Further quantification of continuous drift. (**A**) Number of principal components explaining 50% of variance in normalized spike count matrix for all sessions and all areas. n = 26, 18, and 12 sessions for LEC, MEC, and CA1, respectively. (**B**) Decoding error measured in minutes when decoding 1-min epochs within the session using a linear classifier. Shuffle obtained by shuffling epoch labels. LEC vs MEC: t(42) = -4.70, p = 2.79e^-5^, LEC vs CA1: t(36) = 4.52, p = 6.54e^-5^, MEC vs CA1: t(28) = 0.01, p = 0.99, LEC vs shuffle: t(80) = -27.89, p = 5.65e^-43^, MEC vs shuffle: t(72) = -14.74, p = 1.91e^-23^, CA1 vs shuffle: t(66) = -15.02, p = 5.62e^-23^, two-sample t-test; n = 26, 18, and 12 sessions for LEC, MEC, and CA1, respectively. (**C**) Pairwise comparison of distance traveled in state space between simultaneously recorded brain areas confirms regional differences in drift during identical experiences. Note that in many sessions the distance traveled in one region was correlated with the distance traveled in the other region. Distances calculated in full dimensional space (i.e., population vectors). LEC vs MEC: t(16) = 6.23, p = 1.20e^-5^, LEC vs CA1: t(10) = 2.31, p = 0.04, MEC vs CA1: t(11) = -1.68, p = 0.12, paired t-test; n = 17, 11, and 12 sessions for each region comparison, respectively. (**D**) Example LEC trajectory from rat 26863 showing that drift is immediately present in the animal’s first ever experience foraging in an open field environment. Plotting conventions are the same as in Fig. 1B. (**A-B**) Data represented as mean +/- SEM. (**B-C**) ***p < 0.001, *p < 0.05, ns = not significant.

**Fig. S5.**
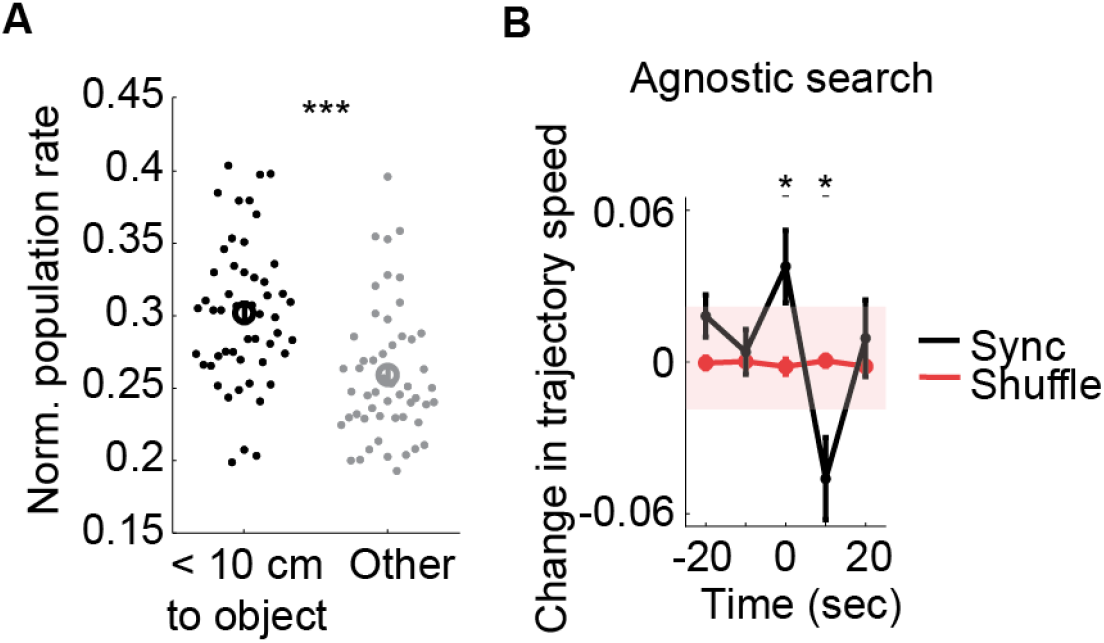
Responses to novel event boundaries. (**A**) Mean population firing rates were higher when animals were near objects (< 10 cm) compared to randomly sampled timepoints not near objects. Summary data from all sessions. Dots represent 10-sec time bins. t(51) = 5.61, p = 8.45e-7, paired t-test, n = 52 timepoints, ***p < 0.001 (**B**) Mean change in trajectory speed for all times of network synchrony (black) during the whole experiment showed time-locked acceleration then deceleration whereas random timepoints (red) did not. Stars mark timepoints where mean is below/above the 1^st^/99^th^ percentile of the shuffled distribution (shaded region). Interaction effect between Sync and Shuffle: F(4, 112) = 4.89, p = 1.10e^-3^, repeated measures ANOVA; n = 15 timepoints. (**A-B**) Data represented as mean +/- SEM.

**Fig. S6.**
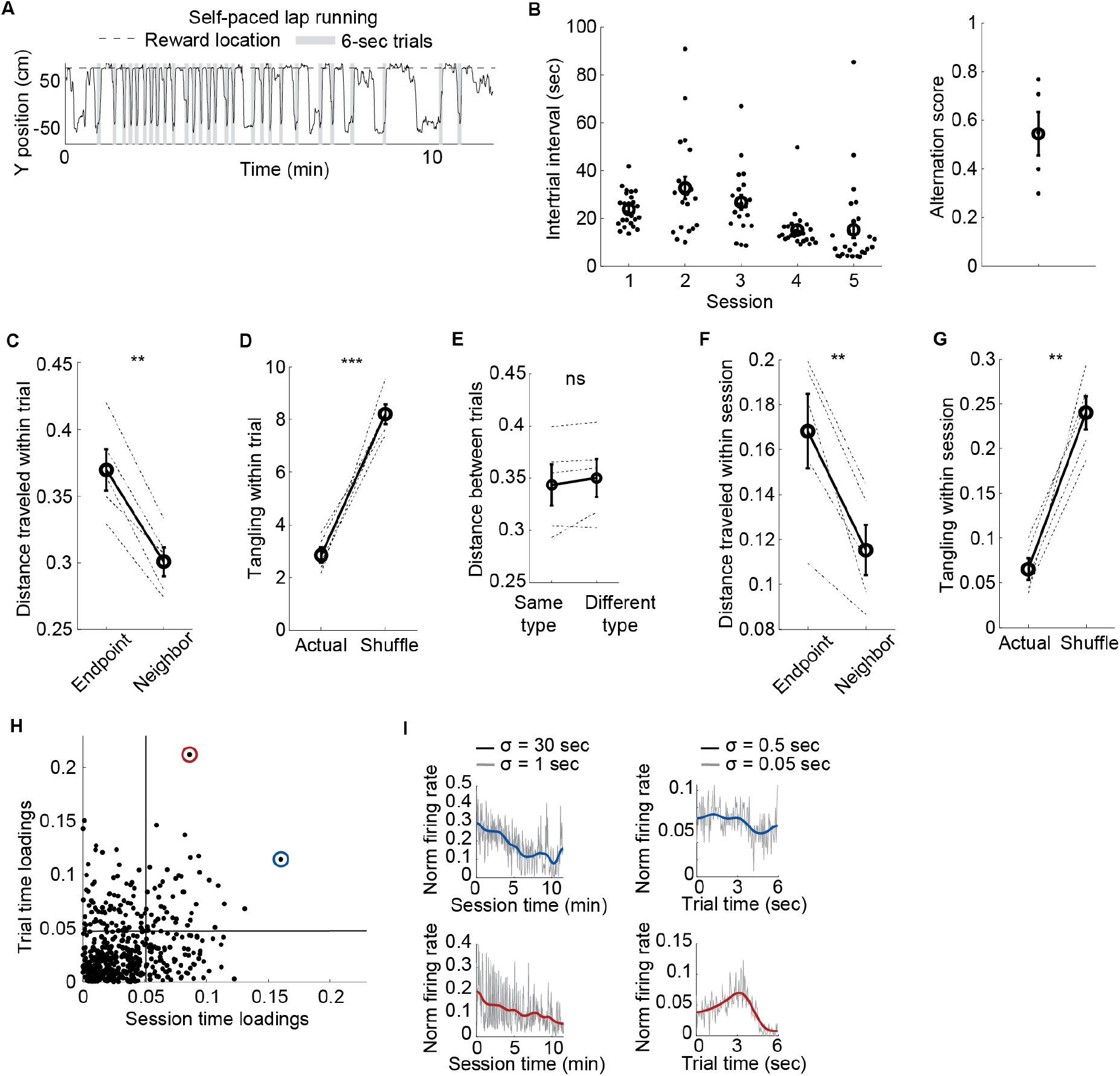
Further quantification of figure-eight data. (**A**) Example behavior during figure-eight task where y-position on the maze over time is shown with a black line, the reward location is marked with a horizontal dashed line, and the 6-sec trials that were extracted post-hoc are shown as shaded gray regions. (**B**) Distribution of intertrial intervals during figure-eight task for each session show that rats ran at their own pace that was variable during a session (left). Dots represent individual intervals. Alternation scores for each session show that rats did not consistently alternate (values near 1) or have bias for one side (values near 0), confirming their free choice running direction on each trial (right). True vs chance value of 0.5: t(4) = 0.51, p = 0.64, one-sample t-test, n = 5 session. (**C**) Distance traveled in full state space was larger between the first and last seconds of the trial compared to neighboring trial times. Endpoint vs neighbor: t(4) = 8.01, p = 1.30e^-3^, paired t-test, n = 5 sessions. (**D**) Trial trajectories showed minimal tangling (LD subspace). Actual vs shuffle: t(4) = -9.17, p = 7.85e^-4^, paired t-test, n = 5 sessions. (**E**) Across all matching trial times, activity was similarly close in the full state space for trials of the same type (left versus right turn trials) vs different type, suggesting that both types were represented by one shared repeating trajectory. Same vs different: t(4) = -1.45, p = 0.22, paired t-test, n = 5 sessions. (**F**) Distance traveled was larger between the first and last minutes of the session compared to neighboring minutes. Endpoint vs neighbor: t(4) = 5.24, p = 6.30e^-3^, paired t-test, n = 5 sessions. (**G**) There was minimal tangling (LD subspace) of neural trajectories during each session. Actual vs shuffle: t(4) = -8.45, p = 1.10e^-3^, paired t-test, n = 5 sessions. (**H**) Multiplexing by individual neurons for session time and trial time was assessed by comparing the (absolute-valued) loadings onto principal components explaining the most variance for either session time or trial time. Vertical and horizontal lines indicate 75^th^ percentiles. Neurons in the top right quadrant multiplexed session and trial time. (**I**) For the circled neurons in (H), firing rates are shown as a function of session time (left) and trial time (right). Traces are smoothed with a Gaussian kernel (width indicated in legend). (**B-G**) Data represented as mean +/- SEM. (**C-G**) ***p < 0.001, **p < 0.01, ns = not significant.

**Fig. S7.**
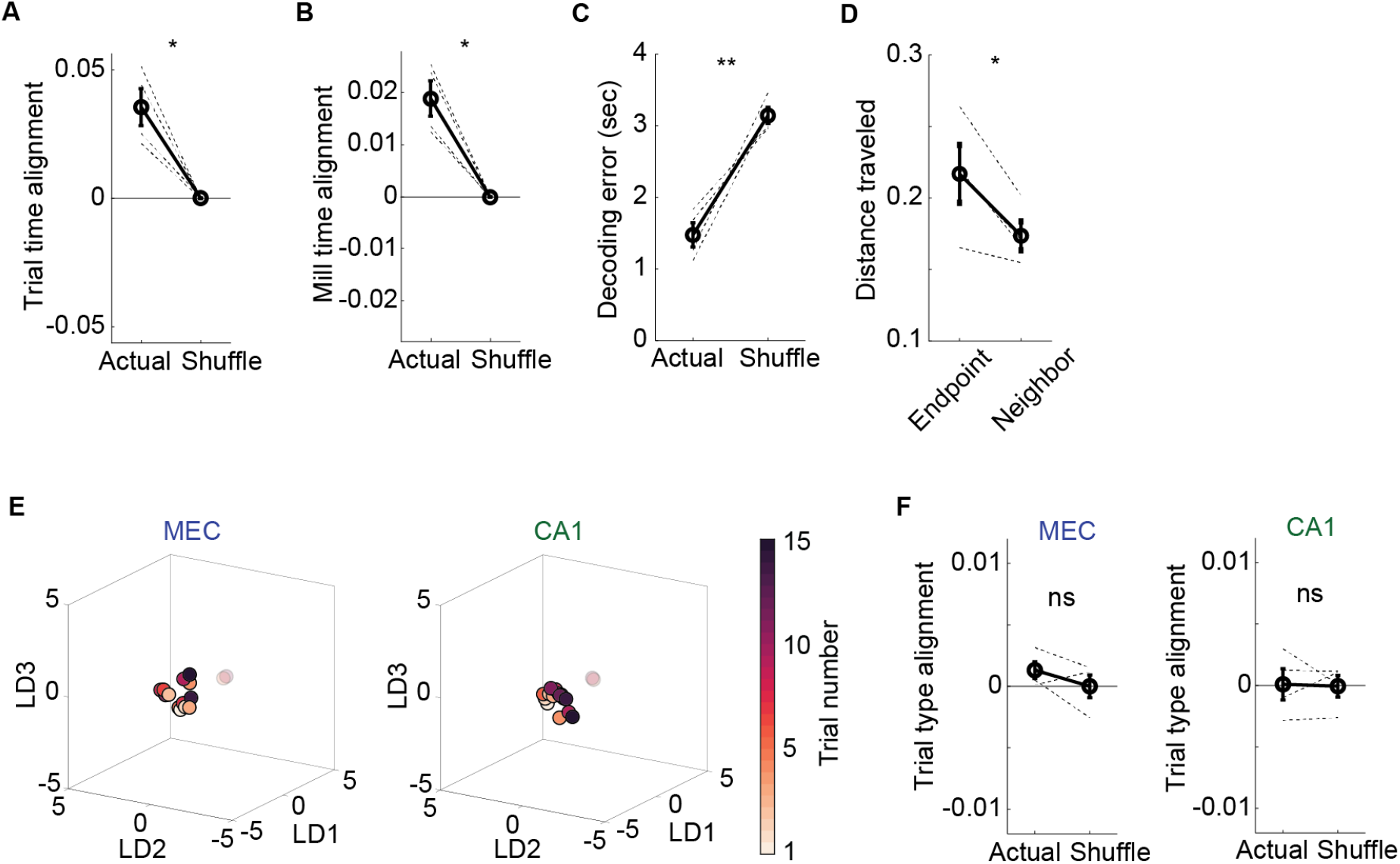
Further quantification of odor sequence data. (**A**) Activity was more similar for matched trial times (i.e., same 1-sec time bin of each trial) compared to mismatched trial times across all trials. Actual vs shuffle: t(3) = 4.88, p = 0.02, paired t-test, n = 4 sessions. (**B**) Activity was more similar for matched mill times compared mismatched mill times across all trials. Actual vs shuffle: t(3) = 5.68, p = 0.01, paired t-test, n = 4 sessions. (**C**) Decoding error for 1-sec mill times using a linear classifier trained on held out trials. Decoding performed on principal components explaining 50% of variance. Actual vs shuffle: t(3) = -6.44, p = 7.58e^-3^, paired t-test; n = 4 sessions. (**D**) Distance traveled was larger between the first and last minutes of the session compared to neighboring minutes. Endpoint vs neighbor: t(3) = 3.81, p = 0.03, paired t-test, n = 4 sessions. (**E**) Example trajectories in LD subspace showing that, in contrast to LEC, sequence runs did not trace parallel trajectories through state space in MEC (left) or CA1 (right). Indeed, there were not clear trajectories in state space in MEC or CA1 compared to the trajectories for LEC presented in Fig. 3J. Dots represent average activity for each trial. Points are colored from light to dark indicating trial number within the session. (**F**) In contrast to LEC, there was no difference in distance between neural activity for matched trial types compared to mismatched trial types across runs for MEC (left) or CA1 (right). MEC vs shuffle: t(3) = 1.53, p = 0.22, paired t-test; CA1 vs shuffle: t(3) = 0.15, p = 0.89, paired t-test; n = 4 sessions. (**A, B, D, F**) Alignment and distance traveled calculated in full n-dimensional state space. (**A-D, F**) Data represented as mean +/- SEM. **p < 0.01, *p < 0.05, ns = not significant.

**Fig. S8.**
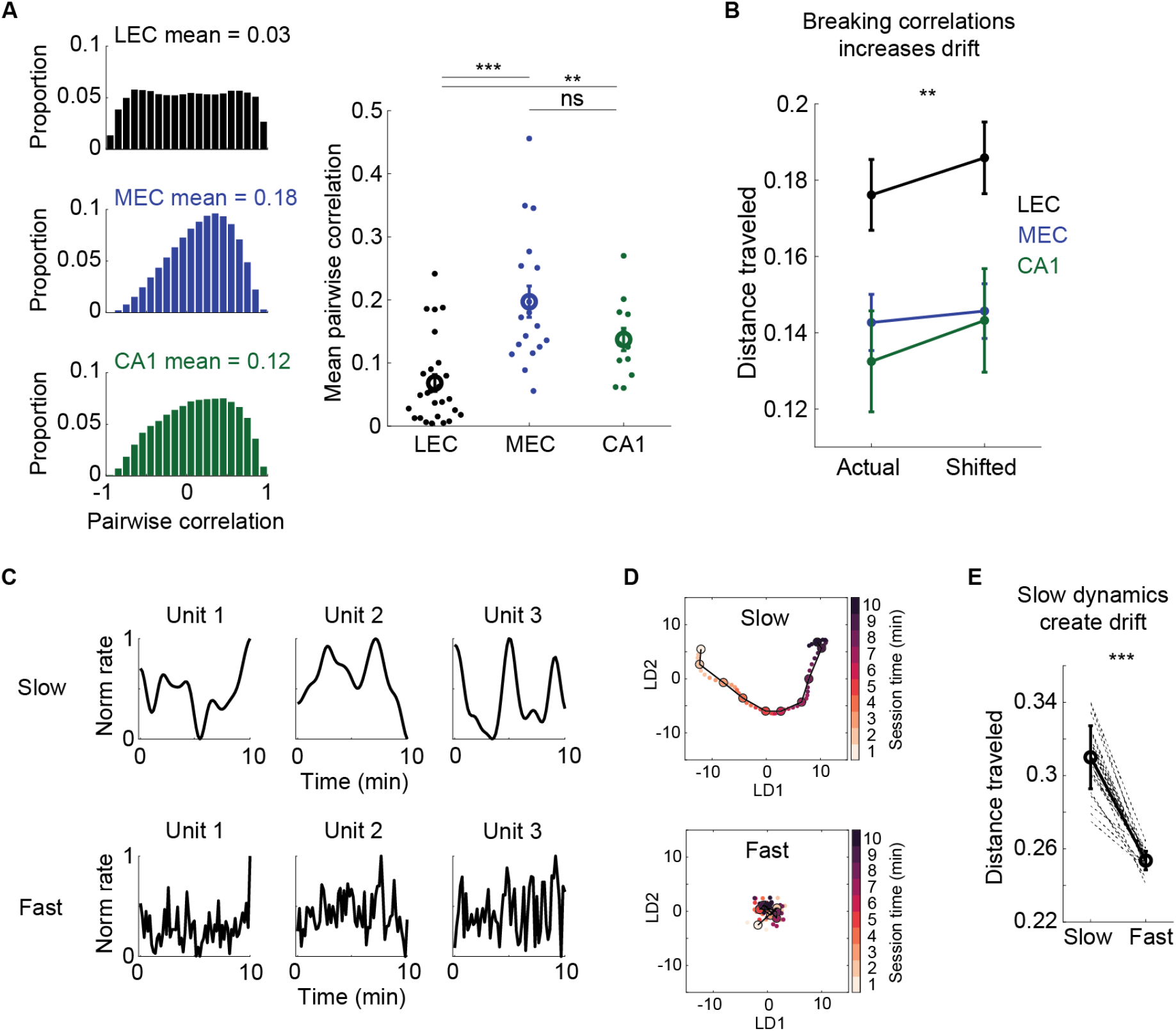
Further quantification of slow dynamics associated with drift. (**A**) Pairwise correlations between neurons using smoothed firing rates (Gaussian width = 30 sec) in an example foraging session (left) shows that LEC has a weaker correlation structure compared to MEC and CA1. Mean pairwise correlation across all neuron pairs within each session (right) shows that LEC correlations across minutes were significantly lower compared to MEC and CA1. LEC vs MEC: t(42) = -4.95, p = 1.26e^-5^, LEC vs CA1: t(36) = -3.00, p = 4.88e^-3^, MEC vs CA1: t(28) = 1.77, p = 0.09; two-sample t-test; n = 26, 18, and 12 sessions for LEC, MEC, and CA1, respectively. (**B**) Breaking correlations within each brain area by independently shifting activity between neurons did not reduce the amount of drift, but rather increased the distance traveled. Main effect between Actual and Shifted: F(1, 53) = 9.41, p = 3.40e^-3^, repeated measures ANOVA; n = 26, 18, and 12 sessions for LEC, MEC, and CA1, respectively (**C-E**) Introducing slow dynamics in simulated, independent Poisson neurons by smoothing their activity with a Gaussian of width 30 sec produces drift compared to those exact same neurons smoothed with a Gaussian of width 1 sec. (**C**) Three simulated units (columns) smoothed with either a 30-sec (top) or 1-sec (bottom) Gaussian kernel. (**D**) Example trajectories showing population drift from slowly varying units (top) and a lack of population drift from quickly varying units (bottom). Simulations of population drift used units as depicted in (C). (**E**)Summary data for 25 simulations showing that distance traveled in state space was significantly greater for slow vs fast dynamics of individual neurons. Slow vs fast: t(24) = 16.95, p = 7.40e^-15^, paired t-test, n = 25 simulations. Thin dashed lines show individual simulations. (**A, B, E**) Data represented as mean +/- SEM. ***p < 0.001, **p < 0.01, ns = not significant.

